# A Synthetic Archaeal Lipid Analog Enables Quantification of Substrate, Products, and Intermediates in the Tetraether Synthase Reaction

**DOI:** 10.64898/2026.09.04.749484

**Authors:** Cody T. Lloyd, Bo Wang, Squire J. Booker

**Affiliations:** Department of Chemistry School of Arts and Sciences, University of Pennsylvania Philadelphia, Pennsylvania, USA; Department of Biochemistry and Biophysics Perelman School of Medicine, University of Pennsylvania Philadelphia, Pennsylvania, USA; Department of Biochemistry and Molecular Biology Pennsylvania State University University Park, Pennsylvania, USA; Department of Chemistry Pennsylvania State University University Park, Pennsylvania, USA; Howard Hughes Medical Institute Chevy Chase, Maryland, USA

**Keywords:** Radical SAM, Tetraether Synthase, Tetraether Lipids, iron-sulfur clusters, S-adenosylmethionine

## Abstract

Glycerol dibiphytanyl glycerol tetraether (GDGT) and macrocyclic diether lipids (mAG) are unique membrane lipids predominantly found in archaea. Both lipids are composed of the same structural components: polar headgroups, glycerol backbones, and ether-linked 40-carbon biphytanyl chains. Tes, a member of the radical *S*-adenosylmethionine enzyme superfamily, synthesizes the biphytanyl chain by catalyzing carbon-carbon (C-C) bond formation between the terminal carbons of two phytanyl chains of the saturated diether lipid substrate. The coupling between the termini of two phytanyl chains on one glycerophospholipid yields mAG, whereas two couplings between the phytanyl termini of two glycerophospholipids yield GDGT. Quantification of the products and presumed intermediates formed during this multistep reaction has been challenging using mass spectrometry due to the lack of appropriate standards for each species. Herein, we designed and synthesized a diether lipid substrate analog containing a fluorescent/UV-visible tag linked to the headgroup, enabling the quantification of each species formed during a Tes reaction. Importantly, under laboratory conditions at 45 °C, we report that GDGT and mAG are initially formed in a 1:1.65 ratio, which differs considerably from the previously assumed ratio.

## Introduction

Archaea inhabit and thrive in some of the harshest environments on Earth, including those characterized by high temperatures, high salinities, and extreme pH levels. The ability to survive in these extreme conditions is primarily attributed to the unique membrane-spanning macrocyclic tetraether lipids that comprise the archaeal membrane (**Fig. 1**).^1-3^ Due to differences in their biosynthetic pathways, archaeal membrane lipids are fundamentally distinct from those found in bacteria and eukarya, a phenomenon known as the “Lipid Divide.” ^4-6^ Archaeal membrane glycerophospholipids exhibit three profound structural differences from those in the other Kingdoms: (1) inversion of stereochemistry at the C2 position of the glycerol backbone, (2) linkage of the alkyl chain to the glycerol backbone via an ether bond, and (3) isoprenoid-based alkyl chains (**Sup. Fig. 1 & Sup. Fig. 2**).^3,4,7^ In archaeal extremophiles, especially the hyperthermophiles, the diether membrane lipids can be modified by linking the terminal carbons of the 20-carbon phytanyl chains to yield the 40-carbon biphytanyl chain, which is observed in the macrocyclic diether lipid (mAG) and glycerol dibiphytanyl glycerol tetraether (GDGT)(**Fig. 1**).^8,9^ The tetraether lipid, GDGT, can be further modified with up to eight cyclopentane rings on the biphytanyl chains (GDGT-1 through GDGT-8), or by forming a bond between two biphytanyl chains of GDGT to give glycerol monoalkyl glycerol tetraether (GMGT)(**Fig. 1**).^10-12^ Several studies have demonstrated that environmental conditions, including temperature, influence the production of tetraether lipids within the membrane.^12,13^ Moreover, recent *in-vivo* work with the archaeal hyperthermophile *Thermococcus kodakarensis* revealed that the presence of tetraether lipids (GDGT and GDGT-1 to -8) in the membrane was essential for the organism’s fitness at higher temperatures.^14^ Their extreme stability, even over geological time scales, and their production at elevated temperatures have made tetraether lipids an ideal ecological proxy for paleoenvironmental temperature reconstruction.^15-17^

**Figure 1.**
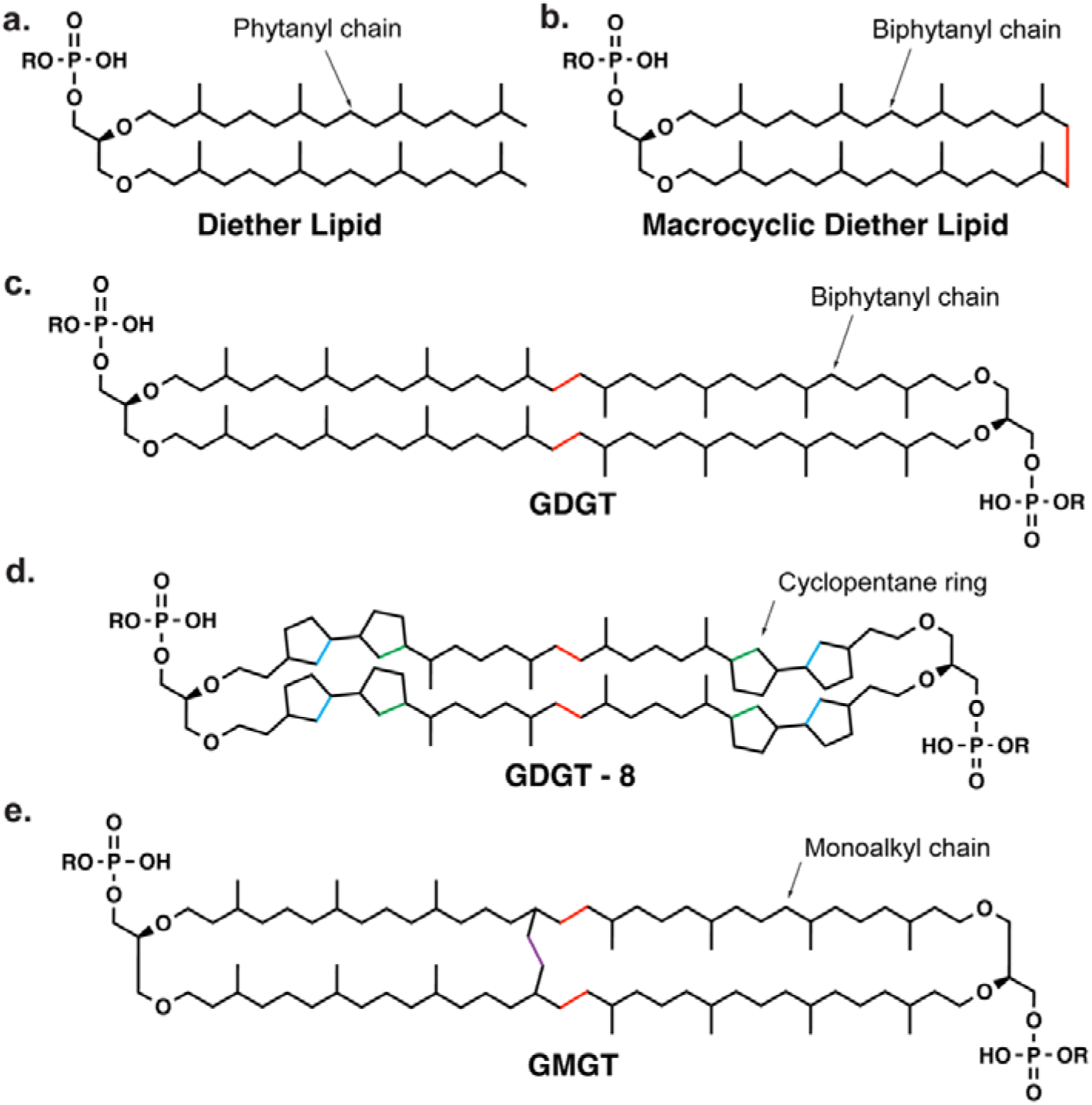
Structures of archaeal membrane lipids: (a.) saturated diether lipid, (b.) macrocyclic diether lipid, (c.) glycerol dibiphytanyl/dialkyl glycerol tetraether with zero cyclopentane rings (GDGT), (d.) GDGT with eight cyclopentane ring modifications (GDGT-8), (e.) glycerol monoalkyl glycerol tetraether (GMGT). The colored C-C bonds in panels b. through e. are modifications catalyzed by Tes (red), GrsA (green), GrsB (blue), and Gms (purple). R corresponds to different headgroups present in naturally occurring archaeal lipids.

The biphytanyl chain observed in all macrocyclic archaeal lipids is formed by tetraether synthase (Tes).^8,9^ During this reaction, Tes couples two completely inert *sp*^*3*^-hybridized carbon (C(sp^3^)) centers. This challenging reaction necessitates radical-dependent chemistry to activate the two carbons undergoing bond formation. Tes is a member of the radical *S*-adenosylmethionine (SAM) enzyme superfamily, using a [Fe_4_S_4_] cluster (denoted [Fe_4_S_4_]_RS_) to catalyze SAM’s reductive cleavage to methionine and a 5′-deoxyadenosyl 5′-radical (5′-dA•). Formation of one C(sp^3^)-C(sp^3^) bond requires two H• abstractions by 5′-dA•, and thus two molecules of SAM, generating the two substrate radicals (**Fig. 2**). The first carbon-centered radical is stabilized by addition to a sulfur atom of a second auxiliary [Fe_4_S_4_] cluster in the C-terminal domain of the protein, denoted [Fe_4_S_4_]_C_. A degraded form of this substrate-cluster intermediate, a thiolated lipid, has been observed by liquid chromatography-mass spectrometry (LC-MS).^8^ The second substrate radical then attacks the carbon of the C-S bond in the substrate-cluster intermediate to form the C(sp^3^)-C(sp^3^) bond observed in the biphytanyl chain.

**Figure 2.**
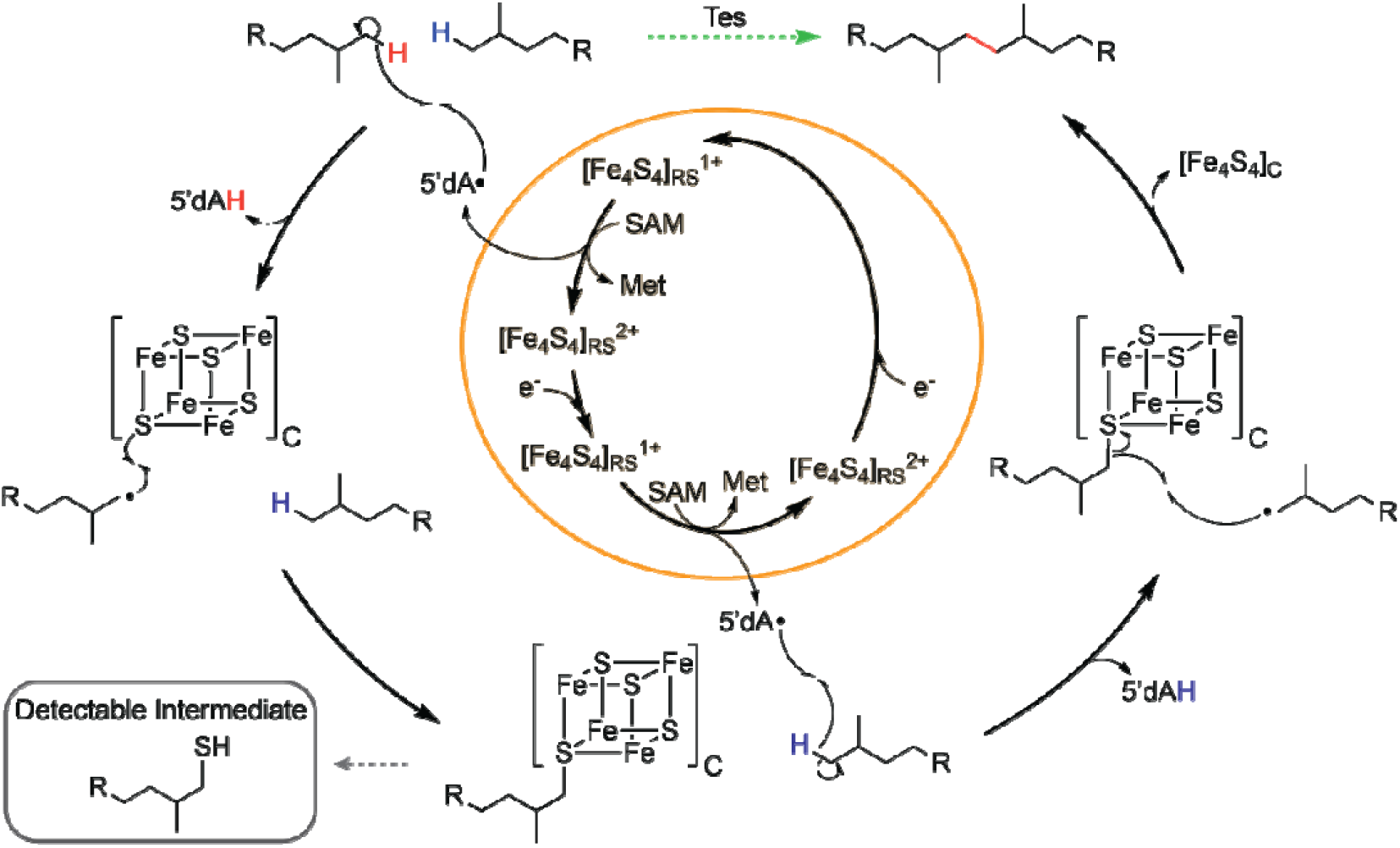
Proposed mechanism for Tes-catalyzed formation of the biphytanyl chain from two phytanyl chains. R denotes the remaining structure of the archaeal lipid (see Fig. 1a). The orange circle indicates the chemistry performed by the [Fe_4_S_4_]_RS_. The dashed green arrow indicates the overall reaction catalyzed by Tes; the newly formed C-C bond is highlighted in red. The dashed gray line indicates a thiolated lipid, which is the LC-MS-detectable, degraded form of the substrate-cluster intermediate that results from destruction of the [Fe_4_S_4_]_C_ cluster during reaction quenching.

The formation of the first C–C bond between phytanyl chains on two different diether lipids is the first committed step toward producing all tetraether lipids. The resulting lipid, glycerol trialkyl glycerol tetraether (GTGT), can undergo a second C–C bond formation by Tes to yield GDGT, the precursor for all subsequent tetraether lipids (**Sup. Fig. 2**). Interestingly, Tes catalyzes the formation of the C–C bonds observed in both tetraether and macrocyclic diether lipids.^8,9^ Promiscuous binding of the four phytanyl chains – two from each diether lipid substrate - in the active site of Tes presumably permits the enzyme to generate an intermolecular product (GDGT) or an intramolecular product (macrocyclic diether).^8^ The resulting macrocyclic diether product is believed to be an insufficient substrate for further modification by Tes due to the binding orientation of the lipid chains in the active site. Therefore, the formation of the macrocyclic diether lipid prevents that lipid from ever becoming a tetraether lipid. Biphytanyl chain formation by Tes is a critical stage in the archaeal lipid biosynthetic pathway and determines the fate of membrane lipids with respect to tetraether lipid production (**Sup. Fig. 2**). Given the profound importance of tetraether lipids in archaeal physiology and paleoenvironmental applications, the factors regulating this partitioning need to be identified.

Membrane lipid profiling studies in some hyperthermophilic archaea have shown that macrocyclic diether lipids are more abundant than tetraether lipids.^18,19^ However, the proportion of tetraether lipids in the membrane increases when the organism is grown at higher temperatures.^13,14^ Nevertheless, the molecular mechanism regulating Tes product partitioning is not understood. In previous studies, products of the Tes reaction were analyzed by mass spectrometry. However, this method is not ideal for quantifying the concentrations of products and intermediates generated, as appropriate standards for each of these species need to be synthesized and quantified. Herein, we engineer an archaeal diether lipid substrate analog containing a fluorescent/UV-vis headgroup, thereby enabling quantification of the substrate, all intermediates, and reaction products. Using this substrate analog, we determine that Tes initially produces GDGT and the macrocyclic diether lipid in a 1:1.65 ratio at 45 °C under laboratory conditions.

## Results and Discussion

### A Novel UV-Handled Archaeal Lipid Substrate Analog

LC-MS was used to identify substrates, intermediates, and products in previous studies of Tes. However, the quantification of these species by this method requires a standard curve prepared from a pure sample of each molecule of interest. Because archaeal lipids are not commercially available, this requires that all of the molecules of interest be chemically synthesized or isolated from a biological source. Although synthetic methods have been developed for a diether lipid with a glycerol headgroup (archaetidylglycerol or AG), a macrocyclic diether lipid, and GDGT, the total synthesis of GTGT and the thiolated intermediates has not been reported.^*20-24*^ Alternatively, these species could be isolated from biological sources (e.g., archaeal organisms or *E. coli* transformed to produce archaeal lipids); however, GTGT and the thiolated intermediates would likely be present at very low abundance.^*25*^ If a single substrate analog containing a fluorescent- or UV-vis-active headgroup is produced, UV-vis or fluorescent spectroscopy could be used to quantify all lipids (substrate, intermediates, and products) relevant to the Tes reaction.

One concern regarding the use of an archaeal lipid substrate analog with a modified headgroup is how the modification could affect substrate binding and enzyme activity. The structure of Tes shows that the lipid headgroup resides in a sizeable solvent-exposed opening and interacts with the protein mainly through water-mediated H-bonds (**Sup. Fig. 3**).^8^ This binding mode enables Tes to synthesize macrocyclic diether and tetraether lipids containing a variety of headgroups. Since the headgroup of the bound lipid is solvent-exposed, we hypothesized that attaching a fluorescent- or UV-vis active species to the headgroup would not negatively affect substrate binding in the enzyme’s active site. Therefore, we synthesized an AG archaeal lipid with a terminal alkyne on the headgroup. Coumarin-343 azide was then ligated using azide-alkyne cycloaddition click chemistry to create a coumarin-343 labeled AG (denoted UV-AG)(**Fig. 3**). Coumarin-343 azide is a fluorophore used in chemical biology that exhibits both fluorescent (λ_*ex*_=437 nm/λ_*em*_=477 nm) and UV-vis (ε=39,000 L⋅mol^−1^ ⋅cm^−1^ ) activity. ^26,27^

**Figure 3.**
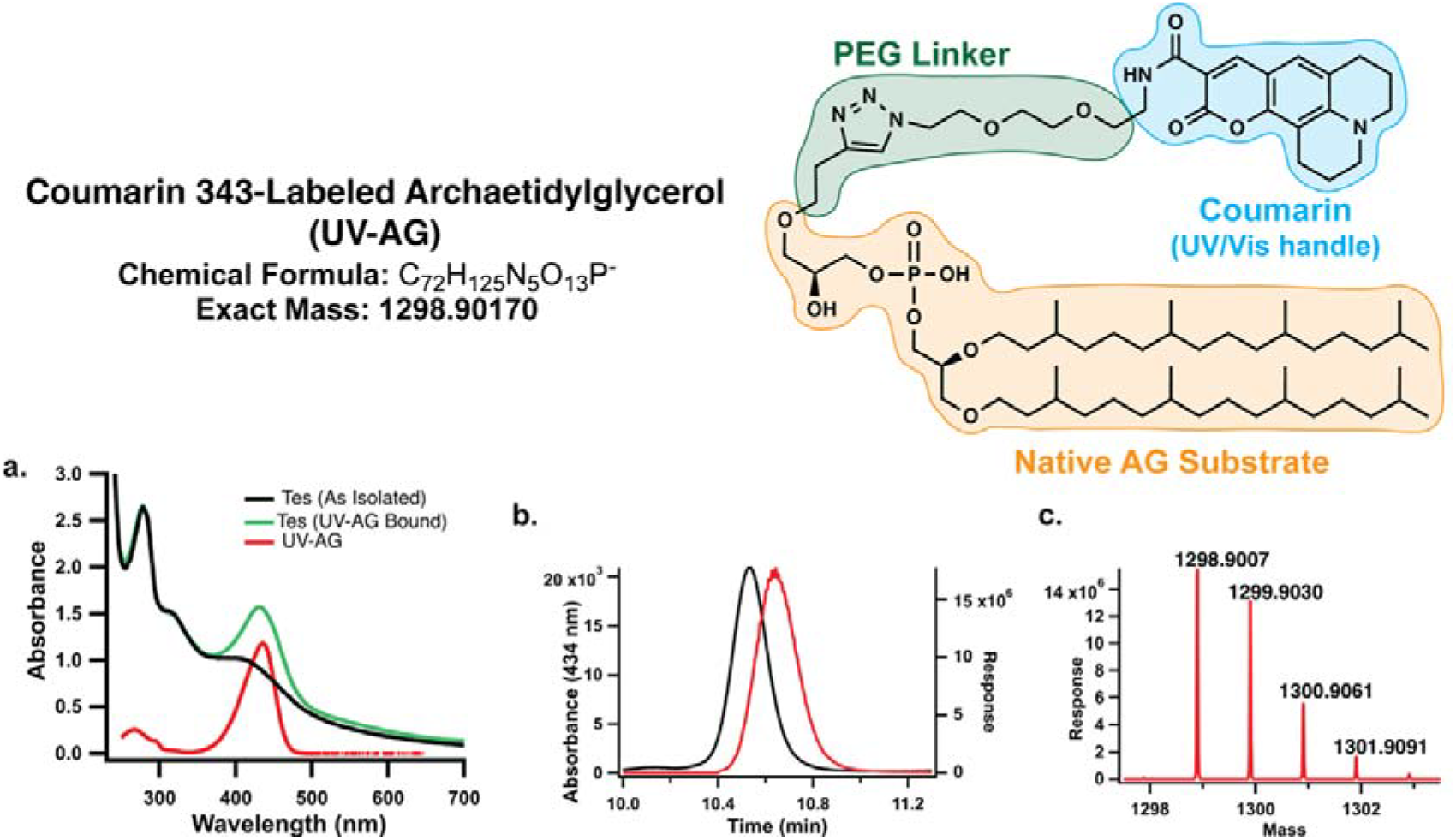
UV-vis and LC-MS characterization of the UV-AG substrate analog. Overall structure of coumarin-343 labeled archaetidylglycerol (UV-AG) with the native AG substrate, PEG linker, and coumarin-343 colored orange, green, and blue, respectively. (a.) UV-vis profile of pure UV-AG (red trace) showing a λ_max_ for the coumarin-343 handle at 434 nm, as-isolated Tes WT (black trace), and Tes WT with UV-AG bound (green trace). (b.) LC-MS spectra for UV-AG by UV-vis (black trace) and MS (red trace) detection. The difference in retention time (RT) between the UV-vis trace (10.54 min) and the MS trace (10.64 min) arises from the transfer tubing connecting the two detectors. (c.) The MS of UV-AG shows the exact mass and the m/z values resulting from natural abundance isotopes.

To assess the binding of UV-AG to Tes, UV-AG was exchanged into the enzyme’s active site, and the UV-AG-bound Tes was analyzed by UV-vis spectroscopy (**Fig. 3a**). Enzymes containing [Fe_4_S_4_] clusters exhibit characteristic UV-vis features around 315 nm and 400 nm, as demonstrated by the UV-vis spectrum of as-isolated Tes (**Fig. 3a**, black trace). UV-AG-bound Tes, however, displays an increased absorbance at 434 nm, the λ_max_ of coumarin-343 (**Fig. 3a**, green trace). These results suggest that the UV-AG lipid binds tightly to Tes. UV-AG-bound Tes was crystallized under anoxic conditions in the presence of 5′-deoxyadenosine (5′dAH) and methionine, and the structure was determined to 2.1 Å resolution (PDB: 37JF, **Fig. 4**). The resulting electron density does not support the modeling of the coumarin headgroup due to its mobility within a large solvent pocket of the crystal lattice. Nevertheless, the UV-AG and AG-bound Tes structures exhibited nearly identical protein architectures, with the C_α_ alignment showing a 0.27 Å RMSD across 453 atoms (**Fig. 4a-c**). Moreover, an evaluation of the active site revealed minimal variations in the positioning of molecules relevant to catalysis, including lipid phytanyl chains, 5′dAH, methionine, and the [Fe_4_S_4_]_RS_ and [Fe_4_S_4_]_C_ clusters (**Fig. 4d-f**). Notably, the positioning of the phytanyl chain terminal carbon relative to the 5′-carbon of 5′dAH is suitable for H• abstraction in both structures, with distances of 3.5 and 3.6 Å. The biochemical and structural characterization of UV-AG-bound Tes supports the use of the lipid analog as a viable substrate for studying the Tes reaction.

**Figure 4.**
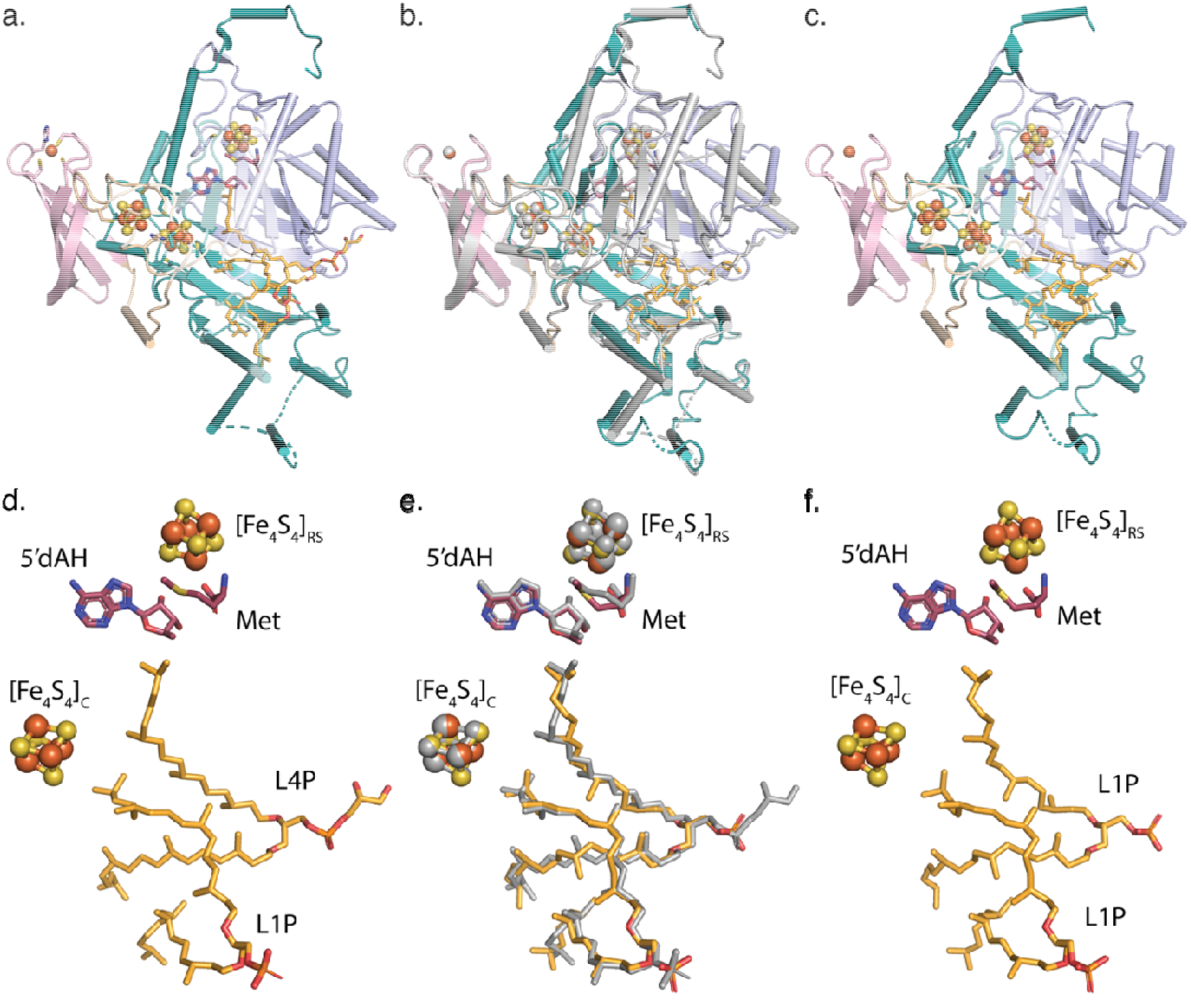
X-ray crystallography structural comparison of Tes with AG or UV-AG bound. Overall structural architecture of Tes with AG bound (a.; PDB 7TOL) and UV-AG bound (c.), showing the rubredoxin domain (light pink), the N-terminal auxiliary cluster domain (wheat), the RS core domain (light blue), and the C-terminal auxiliary cluster (SPASM-like) domain (teal). (b.) Overlay of the AG-bound structur (gray) and UV-AG-bound structure (colors indicated in panel c.) revealing nearly identical protein architectures: Cα alignment of 453 atoms yielded an RMSD of 0.27 Å. (d.-e.) Active site of Tes with AG bound (d.) and UV-AG bound (f.) showing the structural orientation of lipids (orange), 5′dAH (raspberry red), methionine (raspberry red), [Fe_4_ S_4_]_RS_, and [Fe_4_S_4_]_C_.(e.) Active site overlay of the AG-bound structure (gray) and UV-AG-bound structure (colors indicated in panel f.). The active site architectures differ slightly in the positioning of the lipid chains. However, the terminal carbon subject to H-atom abstraction (i.e., the terminal carbon closest to 5′dAH) has the same location.

### Tandem UV/Vis – MS Analysis Enables Lipid Quantification

A preliminary activity assay containing 15 µM UV-AG-bound Tes, 200 µM SAM, and reductant (1 mM titanium citrate) was performed at 45°C to assess whether the lipid substrate, intermediates, and products could be detected by tandem UV-vis-MS. LC UV-vis analysis of the reaction after 5 min revealed six peaks absorbing at 434 nm, consistent with the absorbance of coumarin-343 (**Fig. 5.a**, black trace). Injections of the pure UV-AG lipid identified one of these peaks as the substrate, indicating that Tes produced five additional lipids containing the coumarin-343 headgroup. Subsequent MS analysis of the five additional lipids allowed the assignment as the coumarin-labeled intermediates and products of the Tes reaction based on their exact masses: UV-AG (RT=10.41 min, exact mass=1298.9017), thiolated UV-AG intermediate (UV-AG+S, RT=8.69 min, exact mass=1330.8737), macrocyclic diether product (UV-mAG, RT=10.02 min, exact mass=1296.8861), GTGT (UV-GTGT, RT=19.64 min, exact mass=2596.7950), thiolated GTGT intermediate (UV-GTGT+S, RT=18.25 min, exact mass=2628.7671), and GDGT (UV-GDGT, RT=17.92 min, exact mass=2594.7794)(**Sup. Fig. 4 & Sup. Fig. 5**). Moreover, analysis of this reaction at 30 min showed consumption of the UV-AG substrate and increased production of the UV-mAG and UV-GDGT products (**Fig. 5.b**, red trace). Thus, the LC tandem UV–vis– S method enables the detection of all lipids relevant to the Tes reaction.

**Figure 5.**
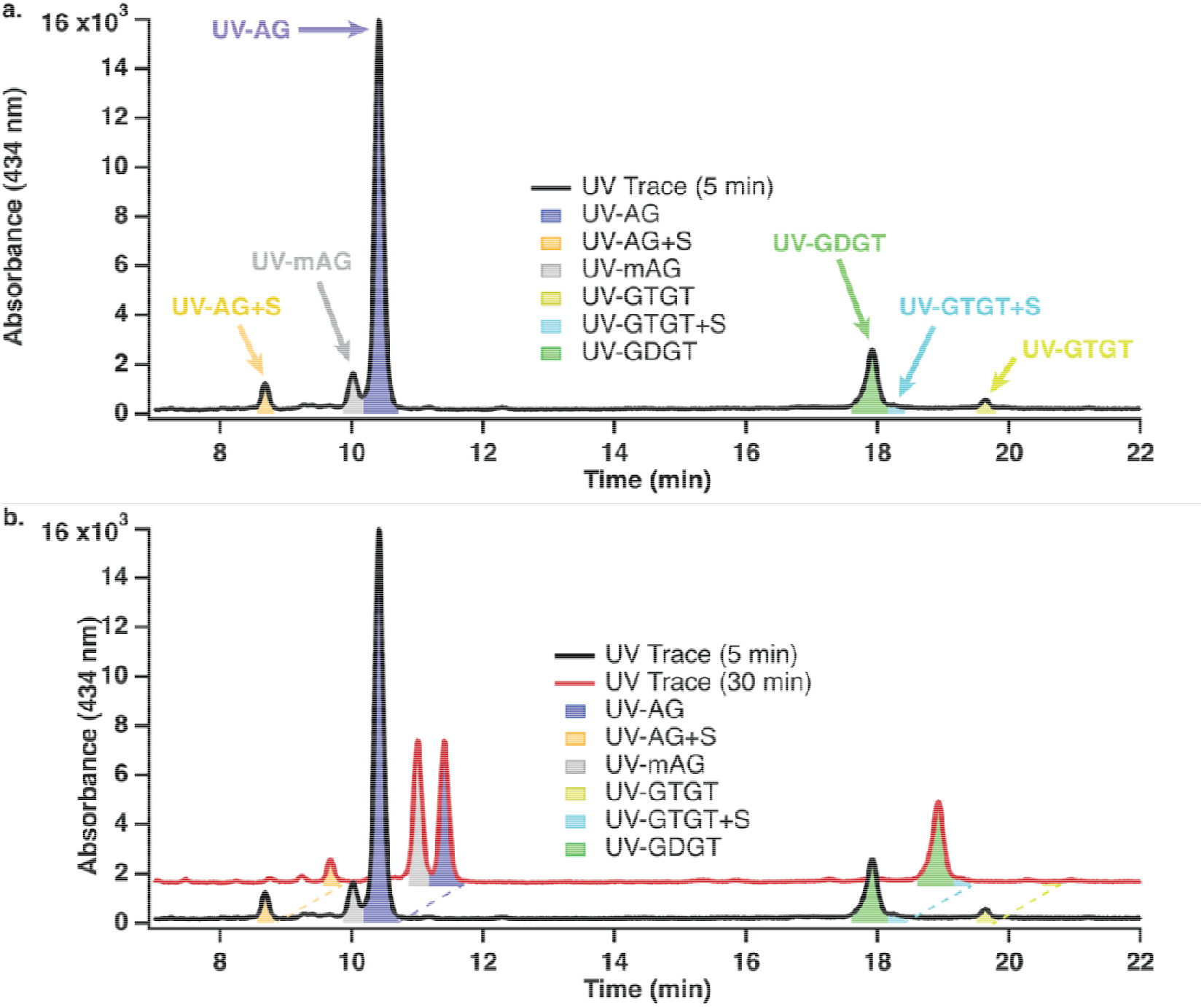
Detection of the UV-handled lipid substrate, intermediates, and products by LC. Chromatographic separation of lipids in the Tes reaction using UV-AG as substrate at 5 min (a.& b., black trace) and 30 min (b., red trace). Detection of the UV-handled lipids at 434 nm enables the identification of all lipids relevant to the reaction: UV-AG+S (RT=8.69 min; orange), UV-mAG (RT=10.02 min; gray), UV-AG (RT=10.41 min; purple), UV-GDGT (RT=17.92 min; green), UV-GTGT+S (RT=18.25 min; blue), and UV-GTGT (RT=19.64 min; yellow).

To quantify the amount of each lipid observed by UV-vis, coumarin-343 azide, the synthetic precursor of UV-AG and possessing the same extinction coefficient as the coumarin handle on UV-AG, was analyzed across a concentration range of 25 nM to 6 µM by the previously described method to generate an external standard (ExSTD) curve(**Fig. 6**). The resulting ExSTD curve was then used to calculate the concentration of coumarin-343 in each peak. Given that the UV-AG substrate has one equivalent of coumarin-343, the resulting UV-AG+S and UV-mAG lipids would also have one equivalent. In contrast, the tetraether lipids (UV-GTGT, UV-GTGT+S, and UV-GDGT) are formed through an intermolecular reaction and would therefore contain two equivalents of the UV-handle. Therefore, if 2 µM of coumarin-343 is observed for a diether lipid peak, that lipid is quantified as 2 µM. In contrast, if 2 µM of coumarin-343 is observed for a tetraether lipid peak, that lipid is quantified as 1 µM. This reasoning was used to quantify UV-AG, UV-AG+S, UV-mAG, UV-GTGT, UV-GTGT+S, and UV-GDGT at each time point of a Tes activity assay. The results showed that, after 15 seconds, 430 nM UV-GDGT and 710 nM UV-mAG formed, reflecting an initial product partitioning ratio of 1:1.65 (**Fig. 6.c**). Throughout the reaction, the steady consumption of UV-AG (blue trace), the formation and decay of the diether intermediate (UV-AG+S, orange trace), and the formation of UV-mAG (gray trace) and UV-GDGT (green trace) products were also observed (**Fig.6.b-c**.). Additionally, the theoretical 5′dAH (i.e., the amount required to yield the quantified lipids via H abstraction) was calculated and compared to the observed 5′dAH (**Fig. 6d**). The time-dependent formation of 5′dAH (blue trace) matched the theoretical 5′dAH (black trace) during the early time points. However, as the reaction progressed, a higher amount of 5′dAH was detected. The detection of excess 5′dAH can be rationalized by the abortive cleavage of SAM, as observed in several other RS enzymes.

**Figure 6.**
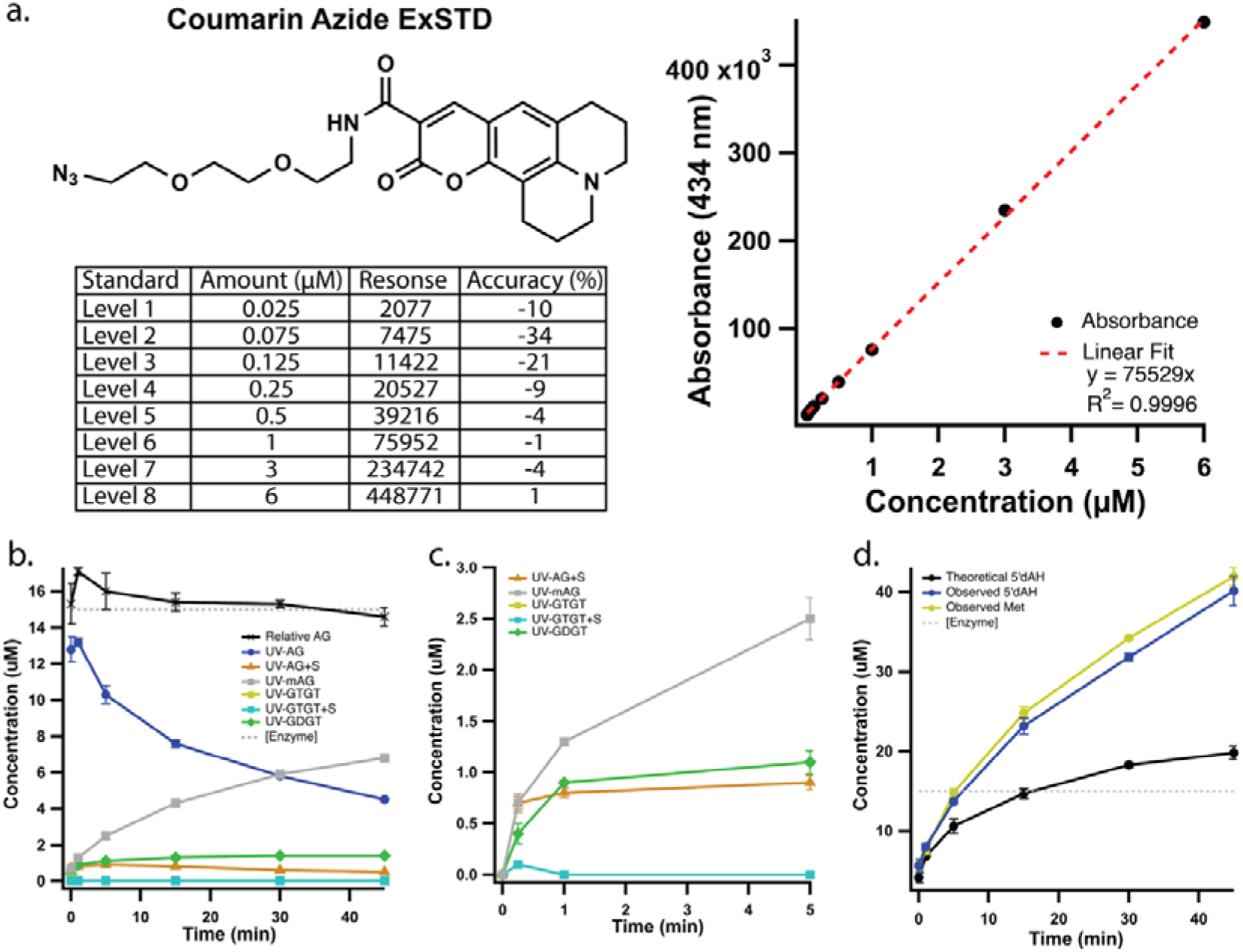
Quantification of the substrate, intermediates, and products using a coumarin-343 azide external standard curve. (a.) Structure of the coumarin azide external standard (top left), table of prepared standard levels and raw data (lower left), and coumarin azide standard curve (right). The accuracies shown in the table reflect the percent difference between the known amount and the amount calculated from the standard curve. (b. & c.) Time-dependent formation of UV lipids during the Tes reaction performed at 45°C showing the consumption of the UV-AG (blue trace) substrate and formation of intermediates and products. The “Relative AG” (black trace) remains constant throughout the reaction, validating the quantification of the lipids. (d.) Time-dependent formation of 5′dAH (blue trace) and methionine (yellow trace) associated with the Tes reaction in panel b. The “theoretical 5′ dAH” (black trace) represents the amount of 5′ dAH required to form the lipids quantified in panel b at each time point. *Assays were performed in triplicate using UV-AG lipid exchanged Tes [15µM], SAM [200 µM], titanium citrate [1 mM], and D-methionine-*d*_3_ [5 µM] as the internal standard. The standard deviation for each timepoint is represented with error bars.

The final validation of this method was to assess the relative amount of coumarin-343 throughout an activity assay. The stable UV-headgroup was the basis for using UV-AG as a substrate analog for Tes. Therefore, the total calculated amount of coumarin-343 should remain unchanged during the reaction. To evaluate the total amount of coumarin-343, the relative amount of UV-AG was calculated from the quantified lipids at each time point. In other words, the amount of substrate required for Tes to generate the intermediates and products was determined and added to the unreacted UV-AG substrate to obtain the relative amount of UV-AG (black trace, **Fig. 6.b**). The relative amount of UV-AG remained constant at approximately 15 µM during a 45-min activity assay using UV-AG-bound Tes. It is worth mentioning that a large amount of bacterial contamination remains bound to Tes following the lipid exchange procedure with UV-AG (**Sup. Fig. 6**). Although the presence of bacterial lipids presents a substantial limitation to our conclusions on product partitioning, particularly the formation of the product resulting from an intermolecular reaction, GDGT, these results suggest that the coumarin-343 azide ExSTD curve accurately quantified all lipids and would be an effective strategy for elucidating product partitioning during the Tes reaction.

## Conclusion

In this work, we developed a strategy to elucidate the partitioning of the macrocyclic archaeal lipid products generated by Tes during *in vitro* activity assays. Using azide-alkyne cycloaddition click chemistry, the fluorescent/UV-visible tag, coumarin-343, was appended to a native archaeal lipid to generate a substrate analog detectable by spectroscopic methods. This strategy is akin to the use of NBD(7-nitrobenz-2-oxa-1,3-diazol-4-yl)-labeled phospholipids for quantifying bacterial lipids and afforded the first quantification of diether and tetraether archaeal lipids using LC-MS instrumentation. In addition to observing and quantifying the substrate, intermediates, and products relevant to the Tes reaction, our results revealed that under single-turnover reaction conditions at 45°C, GDGT and macrocyclic diether lipids are initially formed at a 1:1.65 ratio (**Fig. 6c**). Previous *in vivo* work has shown that environmental conditions, such as temperature and pH, influence macrocyclic lipid composition in archaea.^12-14,28^ However, the molecular mechanisms that regulate this membrane remodeling remain unknown. Although the present work does not address how these environmental factors perturb Tes activity, it provides the first viable strategy to identify the enzymatic factors that regulate this product partitioning *in vitro*. Given the profound importance of macrocyclic lipids, particularly tetraether lipids, in archaeal physiology and their extensive use as biomarkers for paleoenvironmental temperature reconstruction, it is essential to understand how archaea alter their macrocyclic lipid composition.

## Supporting Information

The authors have cited additional references within the Supporting Information. ^[29-39]^

## Acknowledgements

This work was supported by NIH (GM122595 to S.J.B), and the Eberly Family Distinguished Chair in Science (S.J.B.). S.J.B. is an investigator of the Howard Hughes Medical Institute. This research also used the resources of the Berkeley Center for Structural Biology, supported in part by the Howard Hughes Medical Institute. The Advanced Light Source is a Department of Energy Office of Science User Facility under Contract No. DE-AC02-05CH11231. The ALS-ENABLE beamlines are supported in part by the National Institutes of Health, National Institute of General Medical Sciences, grant P30 GM124169.

This article is subject to HHMI’s Open Access to Publications policy. HHMI lab heads have previously granted a nonexclusive CC BY 4.0 license to the public and a sublicensable license to HHMI in their research articles.

Pursuant to those licenses, the author-accepted manuscript of this article can be made freely available under a CC BY 4.0 license immediately upon publication.

## References

1 Valentine, D. L. Adaptations to energy stress dictate the ecology and evolution of the Archaea. Nat Rev Microbiol 5, 316–323 (2007). 10.1038/nrmicro1619

2 Koga, Y. Thermal adaptation of the archaeal and bacterial lipid membranes. Archaea 2012, 789652 (2012). 10.1155/2012/789652

3 Caforio, A. & Driessen, A. J. M. Archaeal phospholipids: Structural properties and biosynthesis. Biochim Biophys Acta Mol Cell Biol Lipids 1862, 1325–1339 (2017). 10.1016/j.bbalip.2016.12.006

4 de Kok, N. A. W. & Driessen, A. J. M. The catalytic and structural basis of archaeal glycerophospholipid biosynthesis. Extremophiles 26, 29 (2022). 10.1007/s00792-022-01277-w

5 Koga, Y. From promiscuity to the lipid divide: on the evolution of distinct membranes in Archaea and Bacteria. J Mol Evol 78, 234–242 (2014). 10.1007/s00239-014-9613-4

6 Woese, C. R., Kandler, O., and Wheelis, M.L. Towards a natural system of organisms: Proposal for the domains Archaea, Bacteria, and Eucarya. Proc Natl Acad Sci U S A 87, 4576–4579 (1990).

7 Koga, Y. & Morii, H. Biosynthesis of ether-type polar lipids in archaea and evolutionary considerations. Microbiol Mol Biol Rev 71, 97–120 (2007). 10.1128/MMBR.00033-06

8 Lloyd, C. T. et al. Discovery, structure and mechanism of a tetraether lipid synthase. Nature 609, 197–203 (2022). 10.1038/s41586-022-05120-2

9 Zeng, Z. et al. Identification of a protein responsible for the synthesis of archaeal membrane-spanning GDGT lipids. Nat Commun 13, 1545 (2022). 10.1038/s41467-022-29264-x

10 Zeng, Z. et al. GDGT cyclization proteins identify the dominant archaeal sources of tetraether lipids in the ocean. Proc Natl Acad Sci U S A 116, 22505–22511 (2019). 10.1073/pnas.1909306116

11 Li, Y. et al. Biosynthesis of GMGT lipids by a radical SAM enzyme associated with anaerobic archaea and oxygen-deficient environments. Nat Commun 15, 5256 (2024). 10.1038/s41467-024-49650-x

12 Garcia, A. A., Chadwick, G. L., Liu, X. L. & Welander, P. V. Identification of two archaeal GDGT lipid-modifying proteins reveals diverse microbes capable of GMGT biosynthesis and modification. Proc Natl Acad Sci U S A 121, e2318761121 (2024). 10.1073/pnas.2318761121

13 Sprott, G. D., Meloche, M. & Richards, J. C. Proportions of diether, macrocyclic diether, and tetraether lipids in Methanococcus jannaschii grown at different temperatures. J Bacteriol 173, 3907–3910 (1991). 10.1128/jb.173.12.3907-3910.1991

14 Liman, G. L. S. et al. Tetraether archaeal lipids promote long-term survival in extreme conditions. Mol Microbiol 121, 882–894 (2024). 10.1111/mmi.15240

15 Wang, J. X., Xie, W., Zhang, Y. G., Meador, T. B. & Zhang, C. L. Evaluating Production of Cyclopentyl Tetraethers by Marine Group II Euryarchaeota in the Pearl River Estuary and Coastal South China Sea: Potential Impact on the TEX86 Paleothermometer. Front Microbiol 8, 2077 (2017). 10.3389/fmicb.2017.02077

16 Ouyang, X., Guo, F. & Bu, H. Lipid biomarkers and pertinent indices from aquatic environment record paleoclimate and paleoenvironment changes. Quaternary Science Reviews 123, 180–192 (2015). 10.1016/j.quascirev.2015.06.029

17 Wang, M., Zheng, Z., Zong, Y., Man, M. & Tian, L. Distributions of soil branched glycerol dialkyl glycerol tetraethers from different climate regions of China. Sci Rep 9, 2761 (2019). 10.1038/s41598-019-39147-9

18 Baumann, L. M. F. et al. Intact polar lipid and core lipid inventory of the hydrothermal vent methanogens Methanocaldococcus villosus and Methanothermococcus okinawensis. Organic Geochemistry 126, 33–42 (2018). 10.1016/j.orggeochem.2018.10.006

19 Comita, P. B., Gagosian, R. B., Pang, H. & Costello, C. E. Structural elucidation of a unique macrocyclic membrane lipid from a new, extremely thermophilic, deep-sea hydrothermal vent archaebacterium, Methanococcus jannaschii. J Biol Chem 259, 15234–15241 (1984).

20 Exterkate, M. et al. A promiscuous archaeal cardiolipin synthase enables construction of diverse natural and unnatural phospholipids. J Biol Chem 296, 100691 (2021). 10.1016/j.jbc.2021.100691

21 Eguchi, T., Arakawa, K., Terachi, T. & Kakinuma, K. Total Synthesis of Archaeal 36-Membered Macrocyclic Diether Lipid. J Org Chem 62, 1924–1933 (1997). 10.1021/jo962327h

22 Eguchi, T., Ibaragi, K. & Kakinuma, K. Total Synthesis of Archaeal 72-Membered Macrocyclic Tetraether Lipids. J Org Chem 63, 2689–2698 (1998). 10.1021/jo972328p

23 Andringa, R. L. H., de Kok, N. A. W., Driessen, A. J. M. & Minnaard, A. J. A Unified Approach for the Total Synthesis of cyclo-Archaeol, iso-Caldarchaeol, Caldarchaeol, and Mycoketide. Angew Chem Int Ed Engl 60, 17497–17503 (2021). 10.1002/anie.202104759

24 Falk, I. D. et al. Enantioselective Total Synthesis of the Archaeal Lipid Parallel GDGT-0 (Isocaldarchaeol)*. Angew Chem Int Ed Engl 60, 17491–17496 (2021). 10.1002/anie.202104051

25 Hoekzema, M., Jiang, J. & Driessen, A. J. M. Optimizing Archaeal Lipid Biosynthesis in Escherichia coli. ACS Synth Biol 13, 2470–2479 (2024). 10.1021/acssynbio.4c00235

26 Teske, N. S., Voigt, J. & Shastri, V. P. Clickable degradable aliphatic polyesters via copolymerization with alkyne epoxy esters: synthesis and postfunctionalization with organic dyes. J Am Chem Soc 136, 10527–10533 (2014). 10.1021/ja505629w

27 Musiol-Kroll, E. M. et al. Polyketide Bioderivatization Using the Promiscuous Acyltransferase KirCII. ACS Synth Biol 6, 421–427 (2017). 10.1021/acssynbio.6b00341

28 Feyhl-Buska, J. et al. Influence of Growth Phase, pH, and Temperature on the Abundance and Composition of Tetraether Lipids in the Thermoacidophile Picrophilus torridus. Front Microbiol 7, 1323 (2016). 10.3389/fmicb.2016.01323

29 Lanz, N. D. et al. RlmN and AtsB as models for the overproduction and characterization of radical SAM proteins. Methods Enzymol 516, 125–152 (2012). 10.1016/B978-0-12-394291-3.00030-7

30 Lanz, N. D. et al. Enhanced Solubilization of Class B Radical S-Adenosylmethionine Methylases by Improved Cobalamin Uptake in Escherichia coli. Biochemistry 57, 1475–1490 (2018). 10.1021/acs.biochem.7b01205

31 Mocniak, L. E., Elkin, K. & Bollinger, J. M., Jr. Lifetimes of the Aglycone Substrates of Specifier Proteins, the Autonomous Iron Enzymes That Dictate the Products of the Glucosinolate-Myrosinase Defense System in Brassica Plants. Biochemistry 59, 2432–2441 (2020). 10.1021/acs.biochem.0c00358

32 Adams, P. D. et al. PHENIX: a comprehensive Python-based system for macromolecular structure solution. Acta Crystallogr D Biol Crystallogr 66, 213–221 (2010). 10.1107/S0907444909052925

33 Bunkoczi, G. et al. Phaser.MRage: automated molecular replacement. Acta Crystallogr D Biol Crystallogr 69, 2276–2286 (2013). 10.1107/S0907444913022750

34 Minor, W., Cymborowski, M., Otwinowski, Z. & Chruszcz, M. HKL-3000: the integration of data reduction and structure solution--from diffraction images to an initial model in minutes. Acta Crystallogr D Biol Crystallogr 62, 859–866 (2006). 10.1107/S0907444906019949

35 Otwinowski, Z. & Minor, W. Processing of X-ray diffraction data collected in oscillation mode. Methods Enzymol 276, 307–326 (1997).

36 Emsley, P., Lohkamp, B., Scott, W. G. & Cowtan, K. Features and development of Coot. Acta Crystallogr D Biol Crystallogr 66, 486–501 (2010). 10.1107/S0907444910007493

37 Smart, O. S., Womack, T. O., Sharff, A., Flensburg, C., Keller, P., & Paciorek, W., Vonrhein, C. and Bricogne, G. Grade, version 1.2.20., <https://www.globalphasing.com.> (2011).

38 Williams, C. J. et al. MolProbity: More and better reference data for improved all-atom structure validation. Protein Sci 27, 293–315 (2018). 10.1002/pro.3330

39 The PyMOL Molecular Graphics Systems v. 2.5 (Schrödinger, 2021).

