## Supporting Information for "A Synthetic Archaeal Lipid Analog Enables Quantification of Substrate, Products, and Intermediates in the Tetraether Synthase Reaction"

### TABLE OF CONTENTS

***Methods & Materials*** **4**

Plasmid construction of Tes WT: 4

Overexpression and purification of Tes: 4

Tes Archaeal Lipid Exchange: 5

Synthesis of UV-AG: 6

Tes Activity Assays: 8

Tes structure determination by X-ray crystallography: 9

#### Methods & Materials

Tetrahydrofuran and dichloromethane were obtained from a JC Meyer solvent dispensing system. SiliaFlash 60 silica gel (230-400 mess) for flash chromatography was purchased from Silicycle Inc. Coumarin 343 azide (**compound 9**) was purchased from Lumiprobe. All other chemicals were of the highest grade available and were purchased from MilliporeSigma.

*The vast majority of the methods used were developed or modified from those previously reported by the Booker Lab in *Lloyd, C.T., et al.*, and are restated below for transparency.[8]

*Plasmid construction of Tes WT:*

The gene encoding Methanocaldococcus jannaschii Tes (gene mj0619, UniProt ID: HMPTM_METJA) was optimized for expression in E. coli and ordered from Invitrogen GeneArt Gene synthesis with an added 5′ NdeI cut site and a 3′ XhoI cut site. The Tes gene was removed from the GeneArt pMA-T vector by digestion with the NdeI and XhoI restriction enzymes and subsequently ligated into linearized pET28a plasmid using T4 DNA ligase. The resulting plasmid was named pMj0619. E. coli DH5α cells were transformed with pMj0619, and the sequence was confirmed by DNA sequencing at Pennsylvania State Genomics Core Facility.

*Overexpression and purification of Tes:*

Expression and purification of Tes WT and variants were modified from previously established methods used to obtain soluble RS enzymes via heterologous expression in E. coli.[29,30] An E. coli BL-21(DE3) strain harboring the pDB1282 and pBAD42-BtuCEDFB plasmids was transformed with the desired plasmid. A single colony of the resulting construct was used to inoculate 200 mL of an LB medium starter culture containing 50 μg/mL kanamycin, 50 μg/mL spectinomycin, and 100 μg/mL ampicillin. The starter culture was incubated overnight at 37°C and shaken at 250 rpm. A 4 mL aliquot of the starter culture was used to inoculate 4 L of ethanolamine minimal medium, containing equivalent antibiotic concentrations, in a non-baffled 6 L erlenmeyer flask and grown at 37°C.[30] At an $OD_{600}$ = 0.6, arabinose was added to the culture to a final concentration of 0.2% (w/v) to induce expression of the isc and btu operons on the pDB1282 and pBAD42-BtuCEDFB, respectively. Simultaneously, 25 μM $FeCl_3$ was added to the medium as the iron source for FeS cluster biogenesis. The cultures were then grown to an $OD_{600}$ = 1.0 and 50 μM IPTG was added to the growth to induce the expression of Tes. The temperature was reduced to 30 °C and the culture was incubated for 5 h before harvesting the cells by centrifugation at 7,000 x g. The harvested cells were flash-frozen and stored at -80°C until protein purification. For reasons that we do not understand, the use of the pBAD42-BtuCEDFB plasmid greatly increased the solubility and yield of Tes. Although the plasmid was generated to enhance the solubility of cobalamin-containing proteins, it has also been found to enhance the solubility of some proteins that do not bind cobalamin.[31]

All remaining steps were performed in a Coy Laboratories anaerobic chamber or in an airtight vessel to ensure an oxygen-free environment. Cell paste was resuspended in lysis buffer (50 mM HEPES, pH 7.5, 300 mM KCl, 10% glycerol, 10 mM β-mercaptoethanol (BME), 4 mM imidazole) for 10 min. The following reagents and enzymes were added to the resulting solution and allowed to incubate for 5 min: 1 mg/mL Lysozyme, 0.1 mg/mL DNAse, 0.17 mg/mL PMSF, 0.8 µg/mL cysteine, 0.7 µg/mL $FeCl_3$, and 0.2% Triton x100. The cell suspension was then sonicated for a total of 5 min (45 seconds on, 7 min off) at 35% amplitude. The lysate was centrifuged at 45,000 x g and the resulting supernatant was loaded onto a Ni-NTA column equilibrated with ~100 mL of Lysis Buffer. The column was washed with 100 mL of lysis buffer to remove all non-His-tagged proteins. Tes was eluted from the column with 75 mL of elution buffer (50 mM HEPES pH 7.5, 300 mM KCl, 10% glycerol, 10 mM BME, 300 mM Imidazole). The eluate was concentrated in a 30 kDa MWCO Amicon Ultra Centrifugal Filter. Tes was buffer exchanged into storage buffer (50 mM HEPES pH 7.5, 300 mM KCl, 20% glycerol, and 1 mM DTT) via a PD-10 desalting column. The resulting protein mixture was further purified by size exclusion chromatography on a HiPrep 26/60 S200 column with an isocratic method using S200 buffer (50 mM HEPES pH 7.5, 300 mM KCl, 10% glycerol, and 10 mM DTT) as mobile phase. Fractions indicative of monomeric Tes were pooled, concentrated, and then buffer exchanged into storage buffer before flash-freezing and storage in liquid nitrogen (LN). The resulting protein is referred to as "as-isolated Tes (Tes AI)."

*Tes Archaeal Lipid Exchange:*

The bacterial phospholipids pulled down during the purification of Tes were exchanged with archaeal lipids by incubation of a 16 mL solution containing 3 µM Tes, 5′dAH at 2.5 mM, methionine at 2.5 mM, and 20 µM synthesized archaeal lipid substrate in storage buffer at 50°C for 1 hr. Following incubation, the solution was exchanged into the storage buffer via a PD-10 desalting column to remove excess lipids. Finally, Tes was concentrated in a 30 kDa MWCO Amicon Ultra Centrifugal Filter. The resulting protein is referred to UV-AG bound Tes.

*Synthesis of UV-AG:*

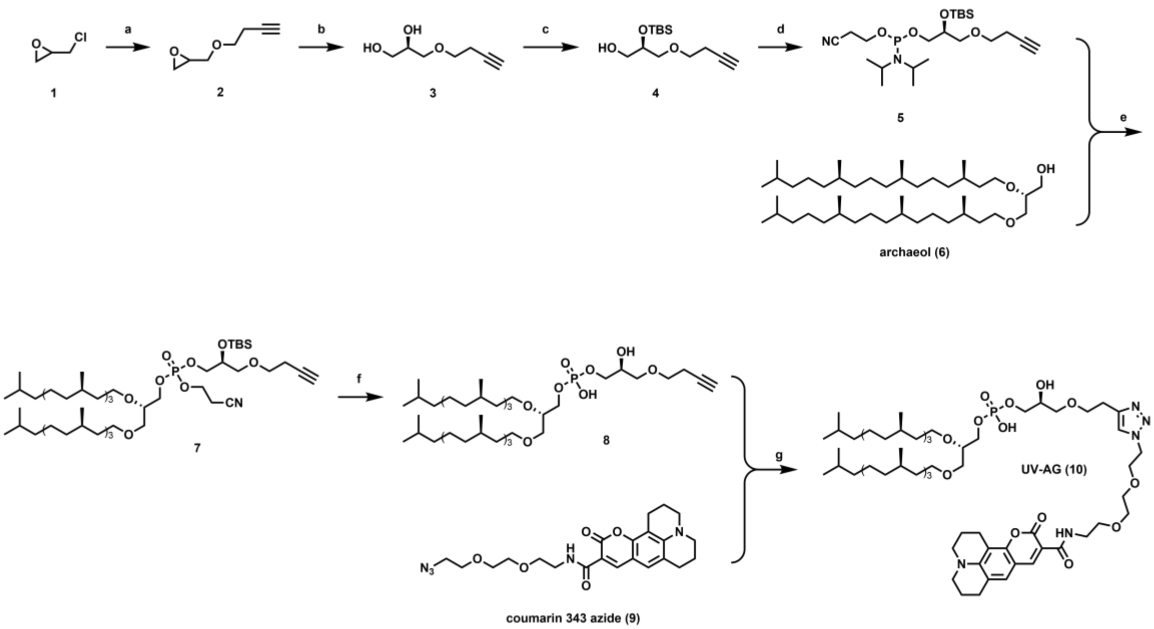

**Conditions and reagents:** **a**) 3-butyn-1-ol, NaOH, tetrabutylammonium hydrogen sulfate, water, rt, overnight, 90%; **b**) (*R,R*)-(−)-*N,N′* -Bis(3,5-di-*tert*-butylsalicylidene)-1,2-cyclohexanediaminocobalt(II), AcOH, $H_2O$, rt, 48h, 44%; c) TBSOTf, TEA, DCM, 0 °C, 1h, then pyridinium *p*-toluenesulfonate, MeOH, 0 °C to rt, 8h, 45% over two steps; **d**) 2-cyanoethyl *N*,*N*-diisopropylchlorophosphoramidite, triethylamine, DMAP, THF, rt, overnight, 90%; e) **6**, tetrazole, DCM, rt, 1h; then $^t$BuOOH, 1h, 80%; f) TBAF, THF, 50 °C, 5h, 77%; g) **9**, $CuSO_4$, ascorbic acid, *t*-butanol, water, 45 °C, 5h, 90%

**Compound 2.** To a stirred solution of sodium hydroxide (25.0 g in 50 mL water) was added epichlorohydrin (33.0 g, 356.8 mmol, 5.0 equiv), 3-butyn-1-ol (5.0 g, 71.3 mmol, 1.0 equiv), and tetrabutylammonium hydrogen sulfate (1.2 g, 3.6 mmol, 0.05 equiv). The resulting solution was stirred at room temperature overnight. After completion, the reaction mixture was extracted by diethyl ether (200 mL). The organic layer was dried over anhydrous sodium sulfate and concentrated *in vacuo*, and the resulting residue was purified by silica gel flash chromatography (hexanes : ethyl acetate = 30:1), giving 8.1 g of compound **2** as colorless oil in a yield of 90%. $^1$H NMR (500 MHz, $CDCl_3$) δ 3.80 (dd, *J* = 11.7, 2.9 Hz, 1H), 3.73 – 3.58 (m, 2H), 3.43 (dd, *J* = 11.7, 5.9 Hz, 1H), 3.21 – 3.10 (m, 1H), 2.80 (t, *J* = 4.6 Hz, 1H), 2.62 (dd, *J* = 5.0, 2.7 Hz, 1H), 2.53 – 2.44 (m, 2H), 2.00 (t, *J* = 2.6 Hz, 1H); $^{13}$C NMR (126 MHz, $CDCl_3$) δ 81.10, 71.60, 69.40, 69.39, 50.74, 44.16, 19.83.

**Compound 3.** To compound **2** (3.2 g, 25.4 mmol, 1.0 equiv) was added (*R*,*R*)-(−)-*N*,*N* ′ -Bis(3,5-di-*tert*-butylsalicylidene)-1,2-cyclohexanediaminocobalt(II) (151.0 mg, 0.25 mmol, 0.01 equiv) and acetic acid (285 μL, 5.0 mmol, 0.2 equiv). The resulting orange solution was stirred at room temperature for 30min. Water (247 μL, 14.0 mmol, 0.55 equiv) was added, and the resulting solution was stirred at room temperature for 48h. After completion, the reaction mixture was purified by silica gel flash chromatography (hexanes : acetone = 1:1), giving 1.6 g of compound **3** as yellowish oil in a yield of 44%. $^1$H NMR (500 MHz, DMSO) δ 4.64 (dd, *J* = 5.0, 1.2 Hz, 1H), 4.53 – 4.44 (m, 1H), 3.59 – 3.52 (m,

1H), 3.52 – 3.45 (m, 2H), 3.43 – 3.39 (m, 1H), 3.34 – 3.28 (m, 3H), 2.83 – 2.75 (m, 1H), 2.42 – 2.35 (m, 2H); $^{13}$C NMR (126 MHz, DMSO) δ 82.45, 72.79, 72.38, 70.98, 69.25, 63.53, 19.66.

**Compound 4.** To an ice-water cooled solution of compound **3** (1.3 g, 9.0 mmol, 1 equiv) and triethylamine (12.5 mL, 90.0 mmol, 10.0 equiv) in DCM (100 mL) was added TBSOTf (6.2 mL, 27.0 mmol, 3.0 equiv) dropwise. The resulting solution was stirred at 0 °C for 1h. After completion, the reaction mixture was diluted with DCM (100 mL) and washed thoroughly with diluted hydrochloric acid, saturated aqueous sodium bicarbonate, water, and brine. The organic layer was dried over anhydrous sodium sulfate and concentrated *in vacuo*, and the resulting residue was used for the next step without further purification.

To an ice-water cooled solution of the resulting residue from previous step in methanol (50 mL) was added PPTS (2.3 g, 9.0 mmol, 1.0 equiv). The resulting solution was gradually warmed up to room temperature and stirred at room temperature for 8h. Then, the reaction mixture was diluted with diethyl ether (200 mL) and washed thoroughly with diluted hydrochloric acid, saturated aqueous sodium bicarbonate, water, and brine. The organic layer was dried over anhydrous sodium sulfate and concentrated *in vacuo*, and the resulting residue was purified by silica gel flash chromatography (hexanes : ethyl acetate = 5:1), giving 1.0 g of compound **4** as colorless oil in a yield of 45% over two steps. $^{1}$H NMR (500 MHz, DMSO) δ 4.58 (t, $J$ = 5.5 Hz, 1H), 3.79 – 3.67 (m, 1H), 3.54 – 3.42 (m, 3H), 3.33 – 3.25 (m, 3H), 2.77 (s, 1H), 2.42 – 2.33 (m, 2H), 0.85 (s, 9H), 0.05 (s, 6H); $^{13}$C NMR (126 MHz, DMSO) δ 82.31, 73.24, 73.12, 72.34, 69.36, 63.50, 26.27, 19.70, 18.40, -4.18, -4.19.

**Compound 5.** To a stirred solution of compound **4** (500 mg, 2.0 mmol, 1.0 equiv), triethylamine (1.4 mL, 10.0 mmol, 5.0 equiv), and DMAP (25 mg, 0.2 mmol, 0.1 equiv) in THF (25 mL) was added 2-cyanoethyl *N*,*N*-diisopropylchlorophosphoramidite (947 mg, 4.0 mmol, 2.0 equiv). The resulting solution was degassed by freeze-vacuum-thaw twice. The reaction was stirred at room temperature overnight. After completion, the reaction mixture was purified by silica gel flash chromatography (hexanes, containing 5% TEA), giving 825 mg of compound **5** as yellowish oil in a yield of 90%. $^{1}$H NMR (500 MHz, $CDCl_3$) δ 3.98 – 3.92 (m, 1H), 3.91 – 3.79 (m, 2H), 3.64 – 3.56 (m, 5H), 3.48 – 3.42 (m, 1H), 2.69 – 2.63 (m, 2H), 2.51 – 2.45 (m, 2H), 1.98 (s, 1H), 1.23 – 1.17 (m, 12H), 0.91 (s, 9H), 0.11 (s, 6H); $^{13}$C NMR (126 MHz, $CDCl_3$) δ 117.65, 81.33, 72.83, 71.47, 69.56, 69.25, 65.01, 58.45, 43.13, 25.83, 24.61, 20.37, 19.82, 18.17, -4.68; $^{31}$P NMR (202 MHz, $CDCl_3$) δ 148.63, 147.97.

**Compound 7.** Archaeol (**6**) was synthesized according to the reported procedure.[32] To a stirred solution of compound **6** (327 mg, 0.5 mmol, 1.0 equiv) and compound **5** (458 mg, 1.0 mmol, 1.0 equiv) in DCM (15 mL) was added tetrazole (2.2 mL, 1.0 mmol, 2.0 equiv, 0.45M solution in acetonitrile). The resulting solution was stirred at room temperature for 1h. *tert*-Butyl hydroperoxide (0.9 mL, 5.0 mmol, 10 equiv, 5.0-6.0M solution in decane) was added to the solution and stirred at room temperature for another 1h. After completion, the reaction mixture was diluted with DCM (100 mL) and washed thoroughly with saturated aqueous sodium thiosulfate, saturated aqueous sodium bicarbonate, water, and brine. The organic layer was dried over anhydrous sodium sulfate and concentrated *in vacuo*, and the resulting residue was purified by silica gel flash chromatography (hexanes : ethyl acetate = 3:1), giving 410 mg of compound **7** as white foam in a yield of 80%. $^{1}$H NMR (500 MHz, $CDCl_3$) δ 4.35 – 3.98 (m, 8H), 3.68 – 3.57 (m, 5H), 3.55 – 3.44 (m, 6H), 2.86 – 2.73 (m, 2H), 2.54 – 2.43 (m, 2H), 2.00 (t, $J$ = 2.6 Hz, 1H), 1.69 – 1.59 (m, 4H), 1.58 – 1.49 (m, 4H), 1.45 – 1.02 (m, 49H), 0.98 – 0.82 (m, 43H), 0.16 – 0.09 (m, 6H); $^{13}$C NMR (126 MHz, $CDCl_3$) δ 116.24, 81.16, 71.93, 71.90, 70.34, 70.22, 69.64, 69.58, 69.44, 69.31, 69.26, 69.03, 67.60, 61.75, 39.39, 37.57, 37.48, 37.44, 37.31, 37.06, 36.62, 32.84, 32.82, 29.96, 29.82, 27.99, 25.76, 24.82, 24.50, 24.40, 22.74, 22.64, 19.82, 19.77, 19.71, 19.69, -4.73; $^{31}$P NMR (202 MHz, $CDCl_3$) δ -1.37, -1.40.

**Compound 8.** To a stirred solution of compound **7** (300 mg, 0.29 mmol, 1.0 equiv) in THF (20 mL) was added TBAF (2.9 mL, 2.9 mmol, 10.0 equiv, 1M solution in THF). The resulting solution was stirred at 50 °C for 5h. After completion, the reaction mixture was diluted with ethyl acetate (100 mL) and washed thoroughly with diluted hydrochloric acid and brine. The organic layer was dried over anhydrous sodium sulfate and concentrated *in vacuo*, and the resulting residue was purified by silica gel flash chromatography (DCM : methanol = 5:1), giving 192 mg of compound **8** as a blackish foam in a yield of 77%. $^{1}$H NMR (500 MHz, $CDCl_3$) δ 4.00 (s, 1H), 3.92 (s, 1H), 3.67 – 3.56 (m, 3H), 3.56 – 3.43 (m, 5H), 2.48 – 2.43 (m, 1H), 1.66 – 1.45 (m, 3H), 1.44 – 0.98 (m, 24H), 0.96 – 0.75 (m, 16H); $^{13}$C NMR (126 MHz, $CDCl_3$) δ 81.26, 71.49, 70.47, 70.20, 69.73, 69.42, 68.88, 50.83, 39.39, 37.66, 37.57, 37.52, 37.33, 37.06, 36.75, 32.90, 32.88, 32.83, 30.05, 29.92, 27.99, 24.82, 24.53, 24.44, 22.74, 22.64, 19.75, 19.73, 19.71, 19.69; $^{31}$P NMR (202 MHz, $CDCl_3$) δ -1.77.

**UV-AG (compound 10).** To a stirred solution of compound **8** (75 mg, 87.3 μmol, 1.0 equiv), compound **9** (46 mg, 104.8 μmol, 1.2 equiv) in a mixed solvent of *t*-butanol (10 mL) and water (10 mL) was added copper sulfate pentahydrate (21.8 mg, 87.3 μmol, 1.0 equiv) and ascorbic acid (30.7 mg, 174.6 μmol, 2.0 equiv). The resulting solution was stirred at 45 °C for 5h. After completion, the reaction mixture was diluted with ethyl acetate (100 mL) and washed thoroughly with water. The organic layer was dried over anhydrous sodium sulfate and concentrated *in vacuo*, and the resulting residue was purified by silica gel flash chromatography (chloroform:methanol:water = 7:3:1), giving 102 mg of UV-AG (**10**) as a green-yellowish foam in a yield of 90%. $^{1}$H NMR (500 MHz, $CDCl_3$) δ 9.36 (s, 1H), 8.86 (s, 1H), 7.77 (s, 1H), 7.14 (s, 1H), 4.54 (s, 2H), 4.27 – 2.60 (m, 48H), 1.96 (s, 4H), 1.61 – 0.96 (m, 69H), 0.94 – 0.76 (m, 41H); $^{13}$C NMR (126 MHz, $CDCl_3$) δ 162.82, 152.66, 148.58, 144.80, 127.95, 127.70, 123.55, 119.78, 108.45, 105.25, 71.45, 70.57, 70.34, 70.04, 69.54, 68.74, 67.35, 65.40, 50.32, 50.18, 49.86, 39.39, 37.67, 37.52, 37.33, 37.10, 36.79, 32.87, 32.83, 30.04, 29.87, 29.72, 27.99, 27.34, 24.82, 24.53, 24.42, 22.75, 22.65, 21.11, 20.18, 20.05, 19.75, 19.64; $^{31}$P NMR (202 MHz, $CDCl_3$) δ -2.78; **HRMS**: calculated for $C_{72}H_{125}N_5O_{13}P^-$ [M-H$^+$]: 1298.9017; found: 1298.9007.

*Tes Activity Assays:*

Activity assays were carried out in triplicate and contained 15 μM lipid exchanged Tes WT, 300 μM SAM, 1 mM TiCitrate, 200 mM KCl, and 10 μM D-methionine-methyl-$d_3$ in 75 mM HEPES, pH 7.5. At each time point, two aliquots were taken from the reaction. For lipid analysis, an aliquot of the reaction was quenched by a five-fold dilution in

IPA:ACN (56.3:43.7 v/v) containing 1 μM phosphatidylglycerol 12:0. For analysis of 5′dAH, an aliquot of the reaction was quenched by a two-fold dilution in 150 mM sulfuric acid.

Quantification of 5′dAH and Met: Reaction aliquots that were quenched in 150 mM sulfuric acid were centrifuged at 13,100 g for 15 min at 4˚C to remove any precipitate. The supernatant was then injected onto an Agilent Technologies 1290 Infinity II series UHPLC system coupled to a 6470 QQQ Agilent Jet Stream electrospray-ionization mass spectrometer. Analytes were chromatographically separated on an Agilent Zorbax Extend-C18 RRHD column (2.1 mm × 50 mm, 1.8 μm particle size) at 32.5˚C that was equilibrated in 95% solvent A (0.1% formic acid, pH 2.6) and 5% solvent B (acetonitrile). Throughout the duration of a single injection, the following gradient was applied: 0 to 0.5 min solvent B was held at 5%, 0.5 to 5 min solvent B increased from 5% to 35%, and from 5 to 6.5 min solvent B increased to 90%. Analytes were detected in positive mode using a multiple-reaction monitoring method. A standard curve of 5′dAH and Met (500 nM through 50 μM) with 5 μM D-methionine-methyl-$d_3$ (internal standard) was prepared for quantification of 5′dAH and Met using the Agilent MassHunter Quantitative Analysis 10.1 Software.

Lipid Quantification: Reaction aliquots that were quenched in IPA:ACN (56.3:43.7 v/v) containing 1 μM phosphatidylglycerol 12:0 were centrifuged at 13,100 g for 15 min at 4˚C to remove any precipitate. The supernatant was then injected onto a Thermo Scientific Vanquish UHPLC system containing a diode array detector coupled to a Thermo Scientific Q Exactive HF-X MS with an H-ESI ion source. Lipids were chromatographically separated on an Agilent Zorbax Extend-C18 column (4.6 mm × 50 mm, 1.8 μm particle size) at 45˚C that was equilibrated in 60% solvent A (60:40 water:ACN with 10 mM ammonium formate and 0.1% formic acid) and 40% solvent B (90:10 isopropanol:ACN with 10 mM ammonium formate and 0.1% formic acid). Throughout the duration of a single injection, the following gradient was applied: 0 to 2 min solvent B increased to75%, from 2 to 11 min solvent B increased to 85%, and from 11 to 17 min solvent B increased to 99%. Analytes were detected by UV-vis at 437 nm. An external standard curve of coumarin azide (25 nM through 6 μM) was prepared for quantification of coumarin-containing lipid substrate, intermediates, and products. For MS, analytes were detected in negative mode using a full-scan method with an H-ESI capillary temperature of 320˚C. From 0 – 12 mins, the full scan MS was collected with an in-source CID of 50.0 eV, a resolution of 120,00, an AGC target of 3e6, and a scan range set to m/z 400-2500. From 12 – 24 mins, the full scan MS was collected with an in-source CID of 30.0 eV, a resolution of 60,00, an AGC target of 1e6, and a scan range set to m/z 1200-2000.

*Tes structure determination by X-ray crystallography:*

General crystallographic methods:

X-ray diffraction datasets were collected at the Berkeley Center for Structural Biology (BCSB) beamlines at the Advanced Light Source at Lawrence Berkeley National Laboratory. All datasets were processed using the HKL2000 or HKL3000 package, and structures were determined by molecular replacement using the program PHASER.[33-36]

Model building and refinement were performed with Coot and phenix.refine, respectively.[33,37] Ligand geometric restraints were obtained from the Grade Web Server (Global Phasing).[38] Structures were validated and analyzed for Ramachandran outliers with the Molprobity server.[39] Figures were prepared using PyMOL.[40]

Crystallization and structure determination of Tes with bound UV-AG:

Brown, plate-shaped crystals of UV-AG bound Tes were generated via hanging drop vapor diffusion method at room temperature in an anaerobic chamber by mixing 1 µL of a solution of UV-AG bound Tes (1 mg/mL) with 1 µL of the well solution [0.1 M HEPES pH 7.5, 30% (*w/v*) PEG 400, 4 mM 5'dAH, and 4 mM methionine]. Crystals were prepared for data collection by mounting on rayon loops followed by soaking in cryoprotectant solution [perfluoropolyether oil (Hampton Research)] and flash-freezing in LN.

The structure was determined by molecular replacement using the coordinates of Tes (PDB 7TOL) as the search model.[8,34,41] Manual model building and refinement were performed in Coot and Phenix, respectively.[33,37] Unmodeled electron density in the active site was assigned as two archaeal lipid molecules (2,3-di-*O*-phytanyl-*sn*-glycero-1-phosphate, L1P).

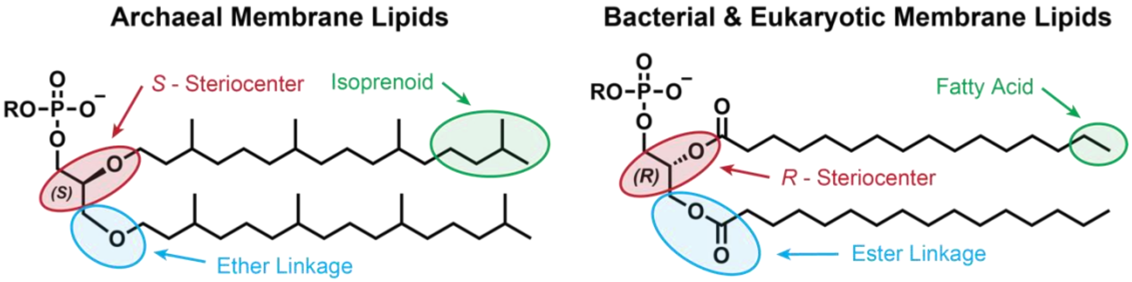

**Supplementary Figure 1**: The lipid divide. Differences between archaeal lipid biosynthesis and bacterial and eukaryotic lipid biosynthesis result in three distinct structural differences: (1) stereochemical inversion of C2 in the glycerol backbone, (2) bond type used to link the alkyl chains to the glycerol backbone, and (3) molecular building blocks for chain elongation.

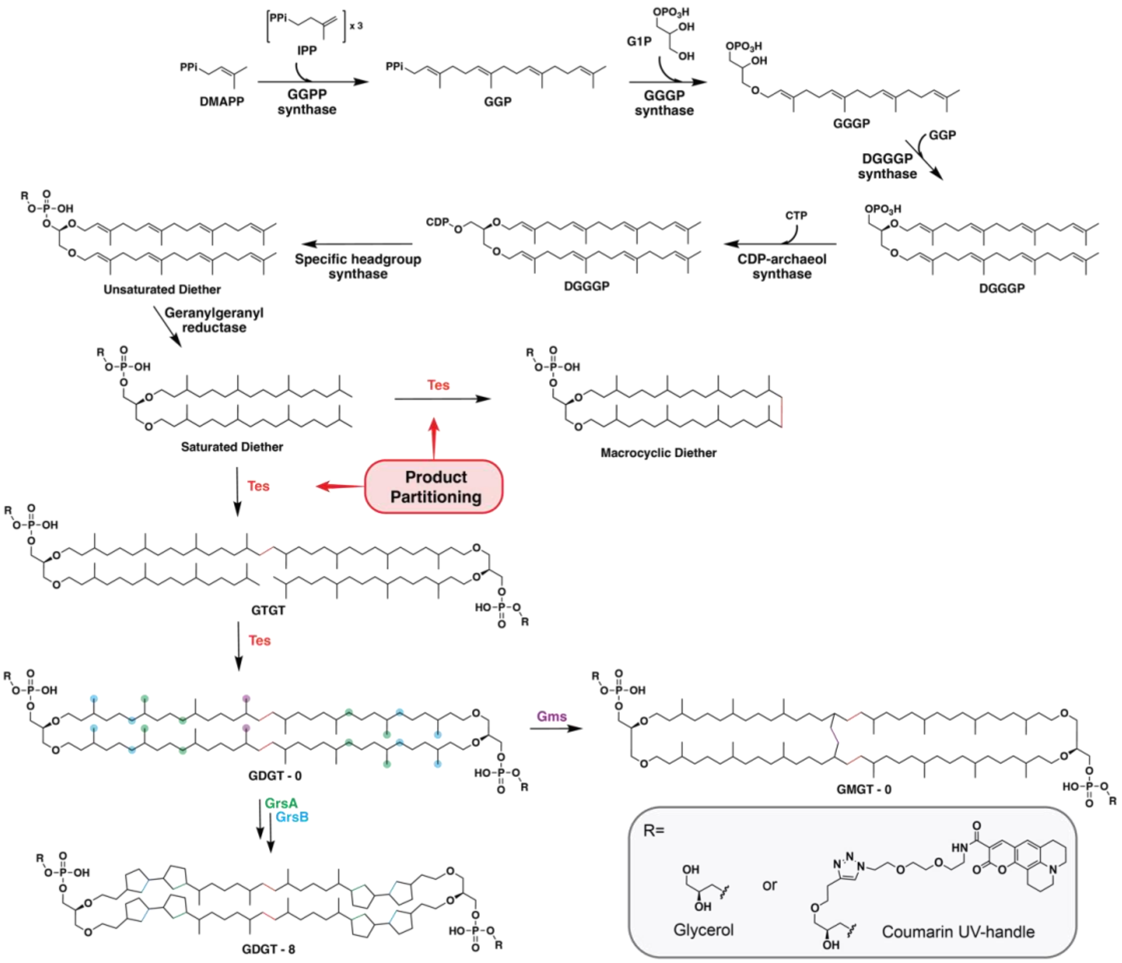

**Supplementary Figure 2**: Archaeal lipid biosynthetic pathway contains a product partition during biphytanyl chain formation. The substrates and enzymes responsible for archaeal lipid biosynthesis are well established. However, biosynthetic access to the tetraether lipids proceeds through a critical product partitioning that results from the reaction promiscuity of Tes. An intramolecular C-C bond formation by Tes results in a macrocyclic diether lipid, which is presumed to be an insufficient substrate for Tes and other downstream tetraether modifying enzymes. Therefore, the intermolecular C-C bond formation resulting in GTGT is the only known route for tetraether lipid formation. Abbreviations: dimethylallyl diphosphate (DMAPP), isopentenyl pyrophosphate (IPP), geranylgeranyl pyrophosphate synthase (GGPP synthase), geranylgeranyl pyrophosphate (GGP), *sn*-glycerol-1-phosphate (G1P), geranylgeranyl glycerol phosphate (GGGP), digeranylgeranyl glycerol phosphate (DGGGP), cytidine diphosphate (CDP), glycerol triialkyl glycerol tetraether (GTGT), Tetraether synthase (Tes), glycerol dibiphytanyl/dialkyl glycerol tetraether (GDGT) with zer0 (-0) through eight (-8) cyclopentane rings, GDGT ring synthase (Grs), glycerol monoalkyl glycerol tetraether (GMGT), GMGT synthase (Gms). *Although several headgroups exist in archaea, R represents glycerol or the Coumarin UV-handle for relevance in this study.

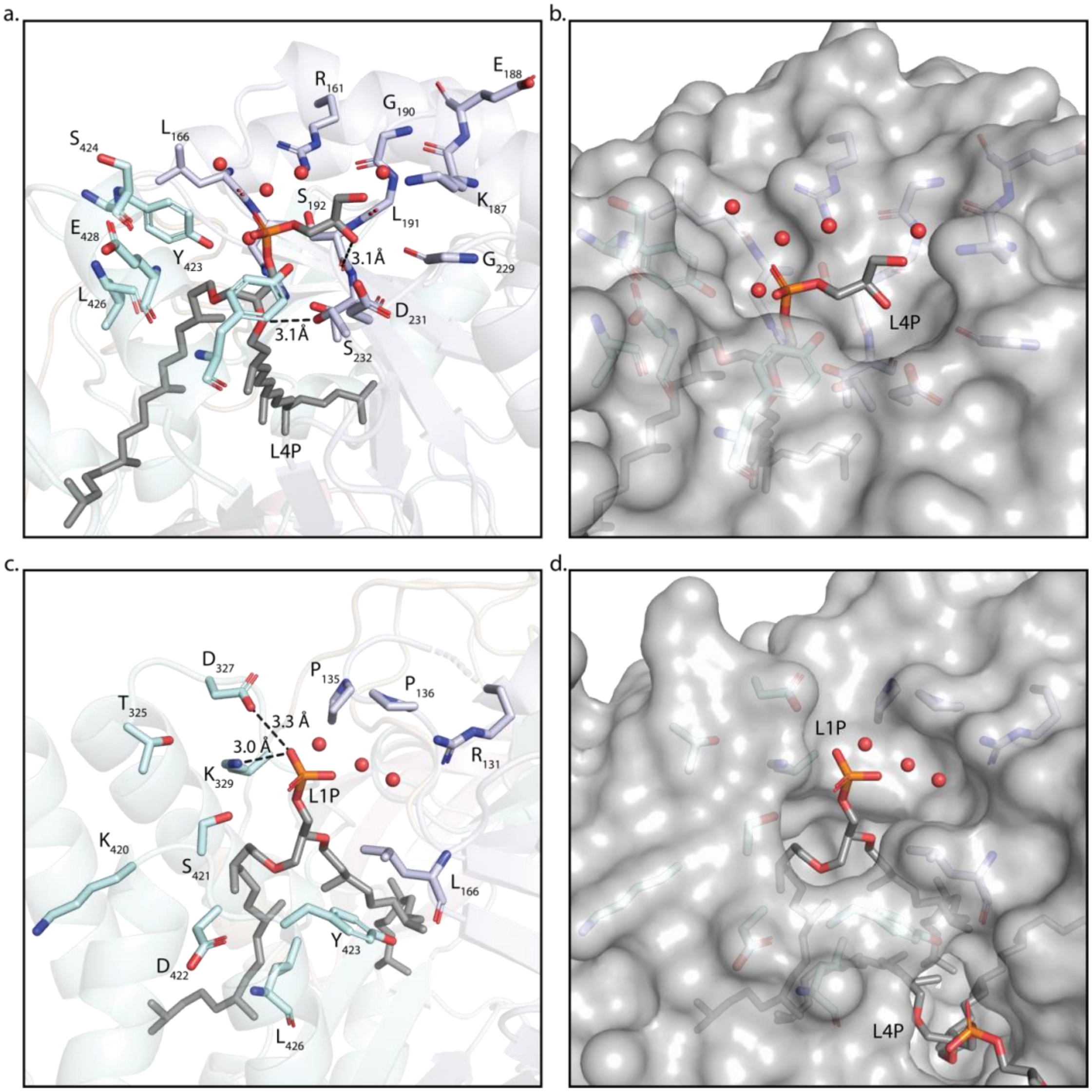

**Supplementary Figure 3**: The nonspecific headgroup binding pocket of Tes.Tes binds the polar headgroup of lipids in a nonspecific fashion, presumably ensuring that all lipids, independent of their headgroup, are sufficient substrates. The lipid binding pocket makes a few direct H-bonds with L1P or L4P (a.&c.). Moreover, only one structurally characterized H-bond interacts with the glycerol headgroup of L4p, while the remaining H-bonds interact with the phosphate and ether linkage. Instead, an H-bonding network with several water molecules mediates the binding of the headgroup to the protein (b.&d.). This large and versatile headgroup binding pocket would allow Tes to accommodate larger polar headgroups, such as the biologically relevant headgroup inositol, and presumable a larger synthetic headgroup - containing a UV-handle - without a substantial impact on the catalytically relevant binding orientation of the phytanyl chains in the active site. *PDB 7TOL was used to make these figures.

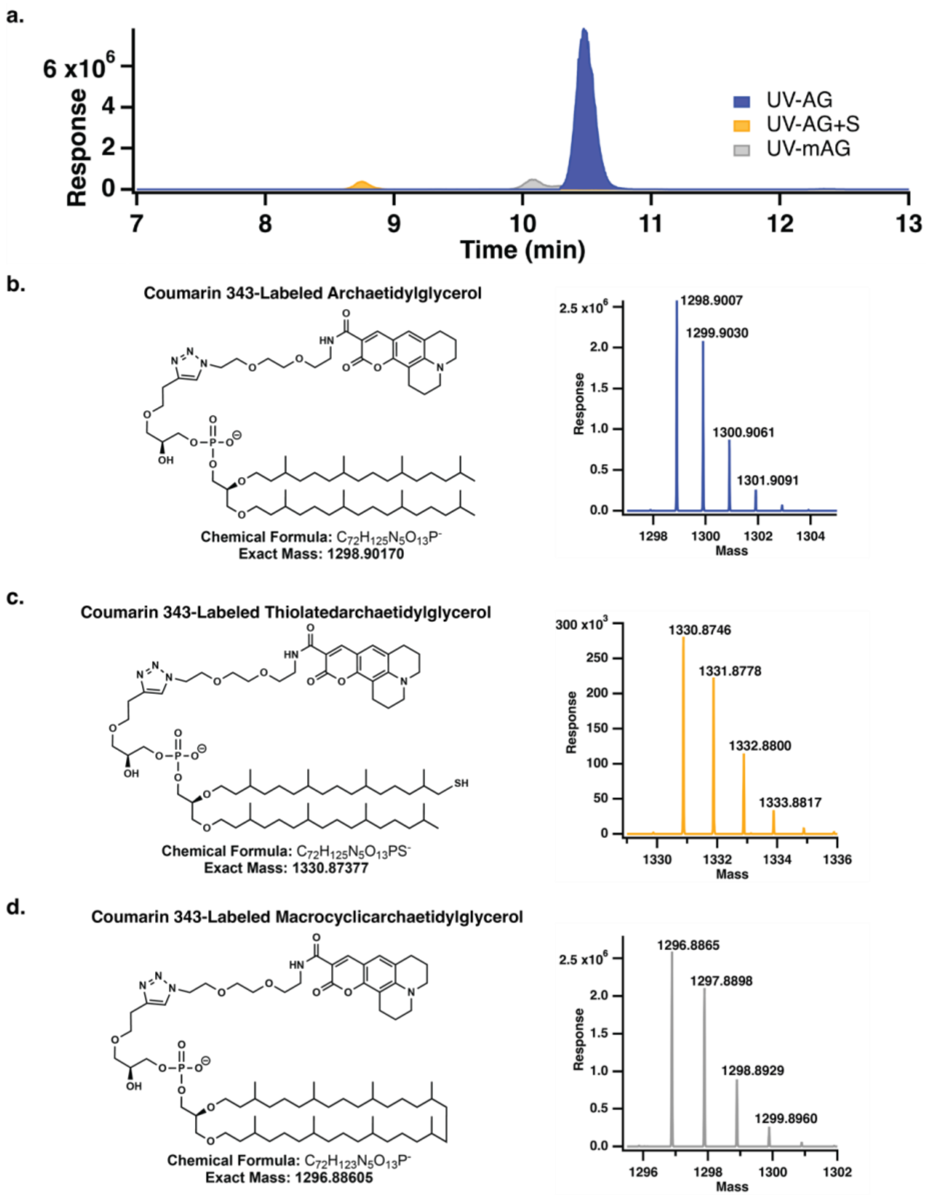

**Supplementary Figure 4**: LC-MS characterization of the diether lipids relevant to the Tes reaction.(a.) LC-MS Extracted ion chromatogram trace (EIC) for Coumarin 343-labeled (UV)-AG (RT=10.48 min; blue trace), UV-AG+S (RT=8.75 min; gray trace), and UV-mAG (RT=10.07 min; orange trace). (b.-d.) Structure of UV-AG (b.), UV-AG+S (c.), and UV-mAG (d.) The right panel shows the mass spectral profile of each lipid's exact mass and the m/z values resulting from natural abundance isotopes, which confirms the molecular formulas.

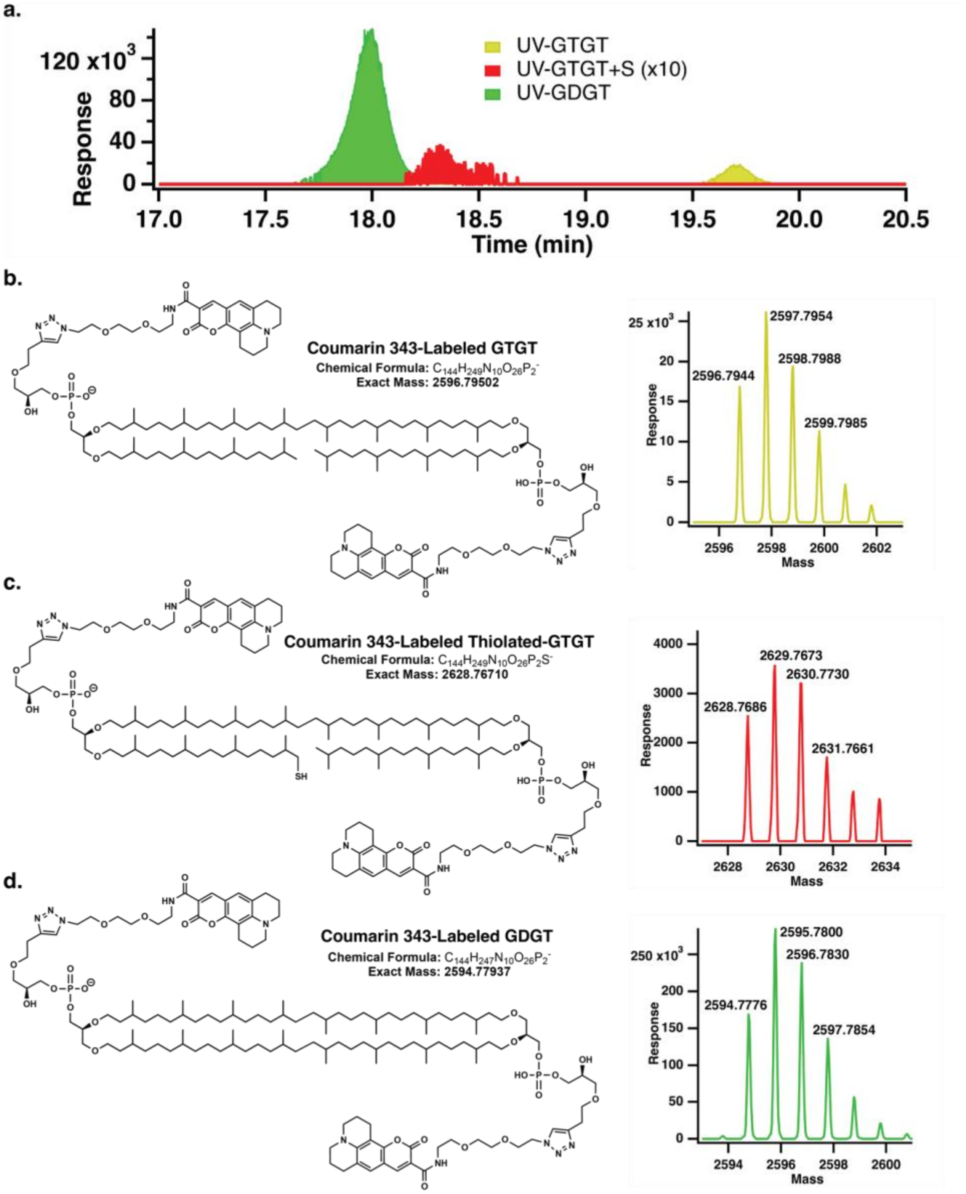

**Supplementary Figure 5**: LC-MS characterization of the tetraether lipids relevant to the Tes reaction.(a.) LC-MS Extracted ion chromatogram trace (EIC) for UV-GTGT (RT=19.72 min; yellow trace), UV-GTGT+S (RT=18.32 min; red trace), and UV-GDGT (RT=18.00 min; green trace). (b.-d.) Structure of UV-GTGT (b.), UV-GTGT+S (c.), and UV-GDGT (d.) The right panel shows the mass spectral profile of each lipid's exact mass and the m/z values resulting from natural abundance isotopes, which confirms the molecular formulas.

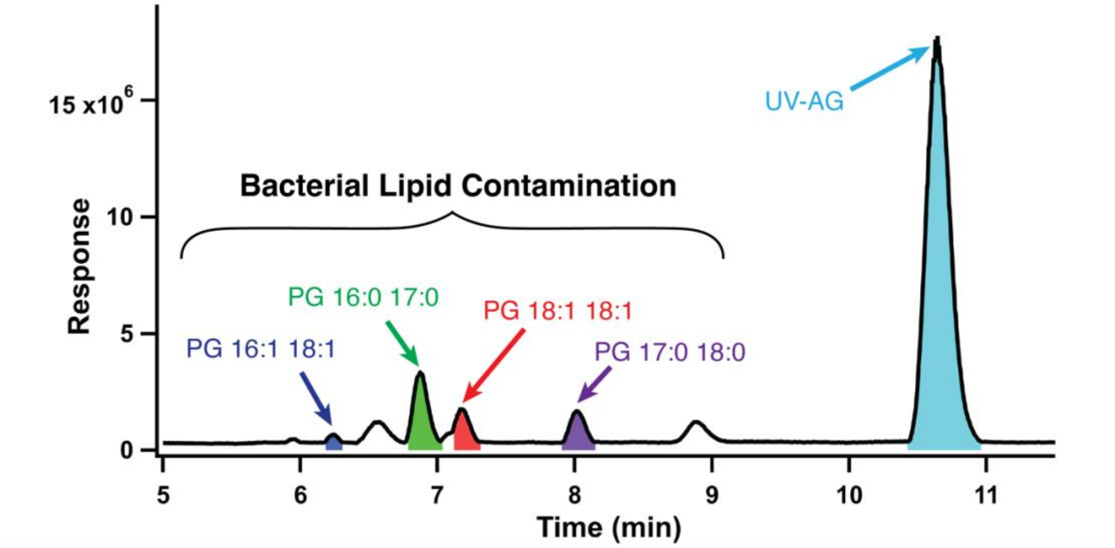

**Supplementary Figure 6**: LC-MS analysis reveals bacterial contamination within the UV-AG LipX Tes. Prior to assays, the bacterial lipids that are advantageously pulled down during protein purification were replaced with the UV-AG lipid during a lipid exchange procedure. However, LC-MS analysis of UV-AG LipX Tes revealed that the protocol did not remove all bacterial lipids. This data suggests that an unknown percentage of Tes is contaminated with one or two bacterial lipids in the active site.

**Supplemental Table 1**. X-ray crystallographic data collection and refinement statistics

| | UV-AG Bound Tes |
|---|---|
| **Data collection** | |
| Space group | $P\ 2_12_12_1$ |
| Wavelength (Å) | 1.00013 |
| Cell dimensions | |
| *a*, *b*, *c* (Å) | 56.28, 74.03, 113.98 |
| *α, β, γ* (°) | 90, 90, 90 |
| Resolution (Å) | 45.16 – 1.97 (2.02 -1.97) |
| No. of unique reflections | 31,406 |
| $R_{sym}$ or $R_{\mathrm{merge}}$ | 0.06 (1.49) |
| $R_{\mathrm{pim}}$ | 0.03 (0.74) |
| $I$ / σ$I$ | 11.4 (0.7) |
| $CC_{1/2}$ | 0.999 (0.420) |
| Completeness (%) | 92.2 (94.0) |
| Redundancy | 4.1 (4.2) |
| **Refinement** | |
| Resolution (Å) | 44.81 – 1.97 (2.02 – 1.97) |
| No. reflections | 31,320 |
| $R_{\mathrm{work}}$ / $R_{\mathrm{free}}$ | 0.2.001 / 0.2440 |
| No. atoms | 4154 |
| Protein | 3962 |
| Ligand/ion | 143 |
| Water | 49 |
| *B*-factors Å$^2$ | |
| Protein | 64.96 |
| Ligand/ion | 64.32 |
| Water | 50.83 |
| R.m.s. deviations | |
| Bond lengths (Å) | 0.010 |
| Bond angles (°) | 1.46 |
| Clashscore | 11.89 |
| Ramachandran | |
| Most favored (%) | 97.76 |
| Allowed (%) | 2.24 |
| Outliers (%) | 0 |
| Number of TLS groups | 4 |
| PDB accession code | 37JF |

*Values in parentheses are for the highest-resolution shell. All structures result from a single crystal.

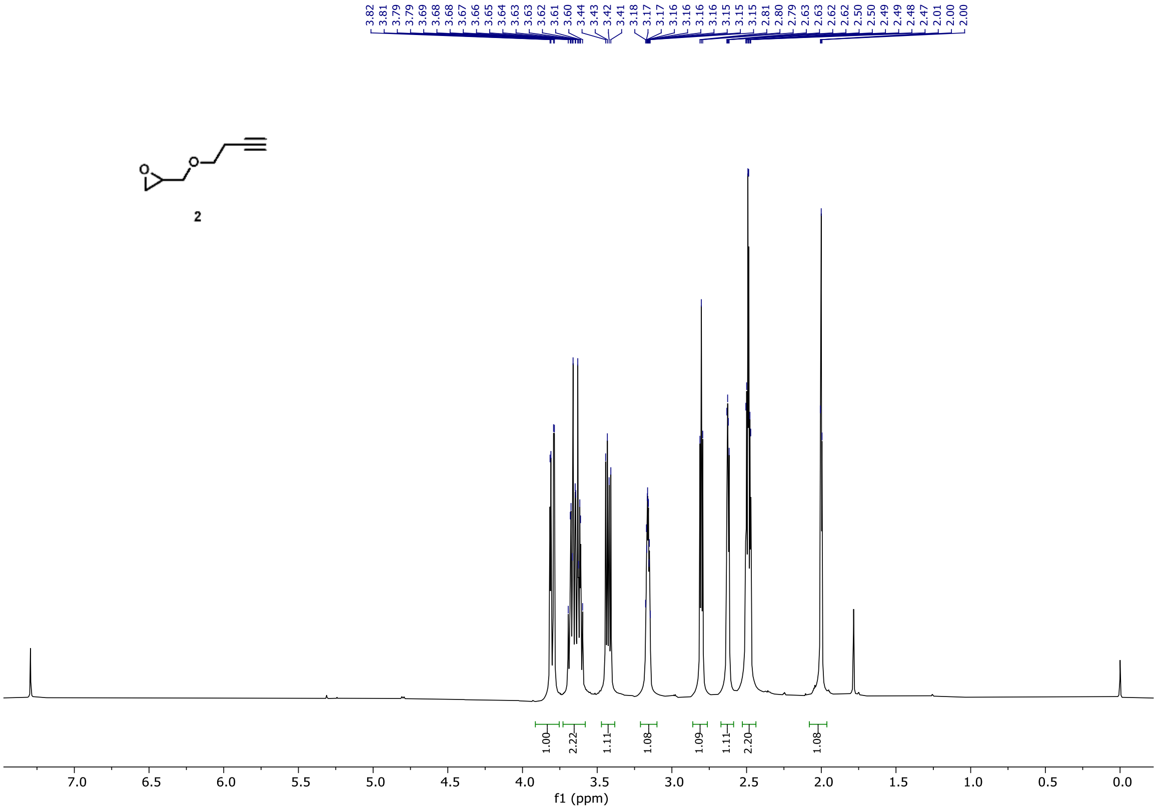

**Supplementary Figure 7**: $^1$H NMR Spectrum of Compound 2.

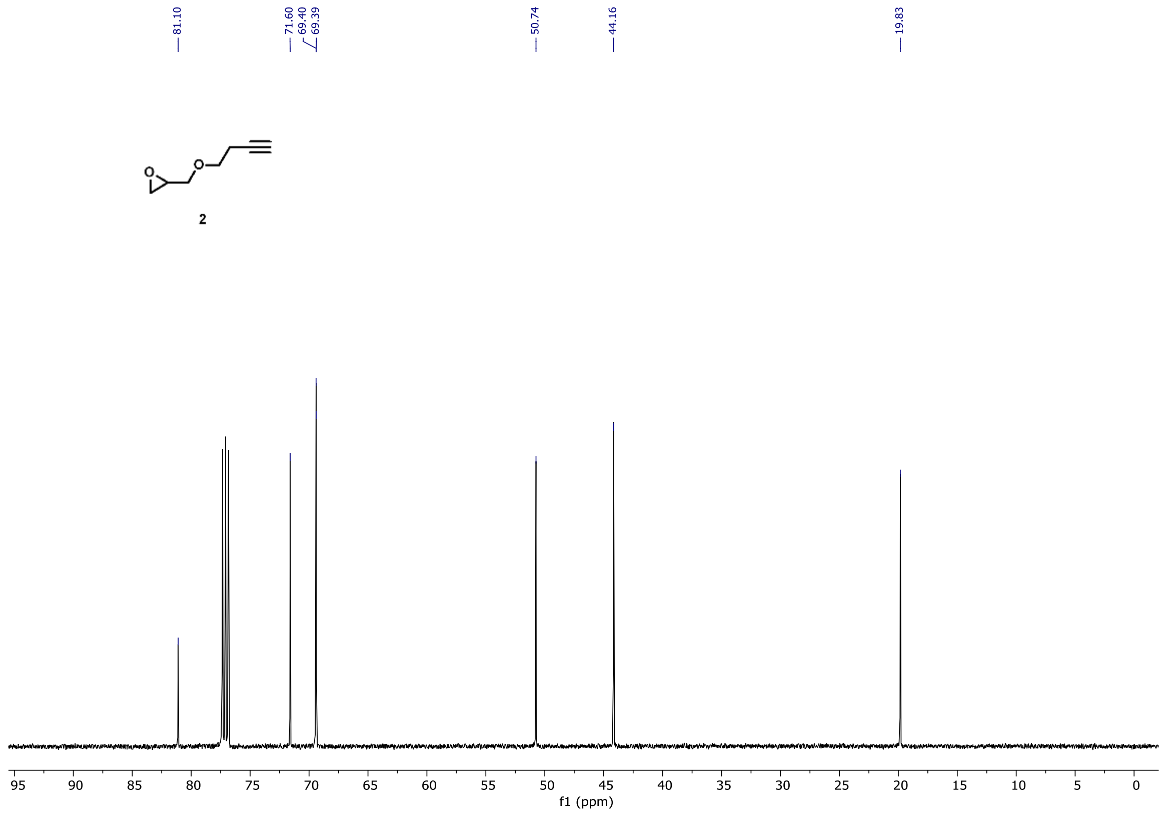

**Supplementary Figure 8**: $^{13}C$ NMR Spectrum of Compound 2.

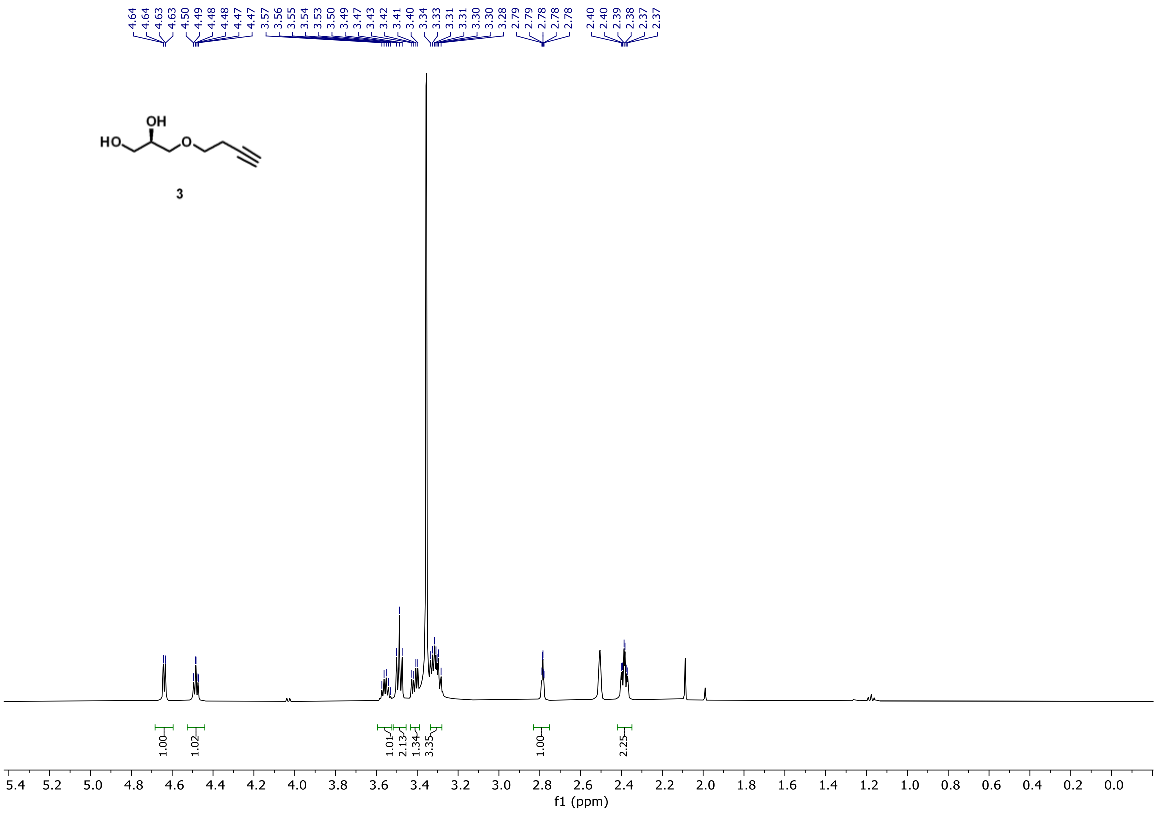

**Supplementary Figure 9**: $^{1}$H NMR Spectrum of Compound 3.

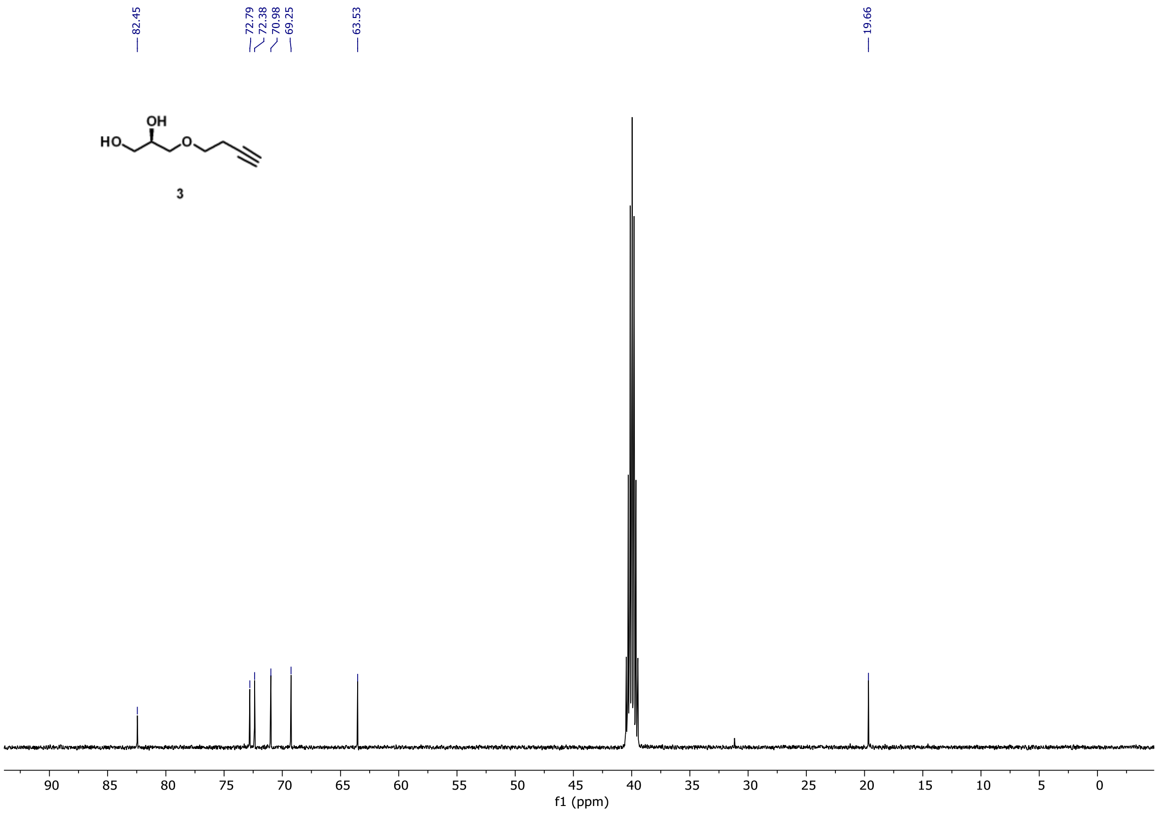

**Supplementary Figure 10**: $^{13}C$ NMR Spectrum of Compound 3.

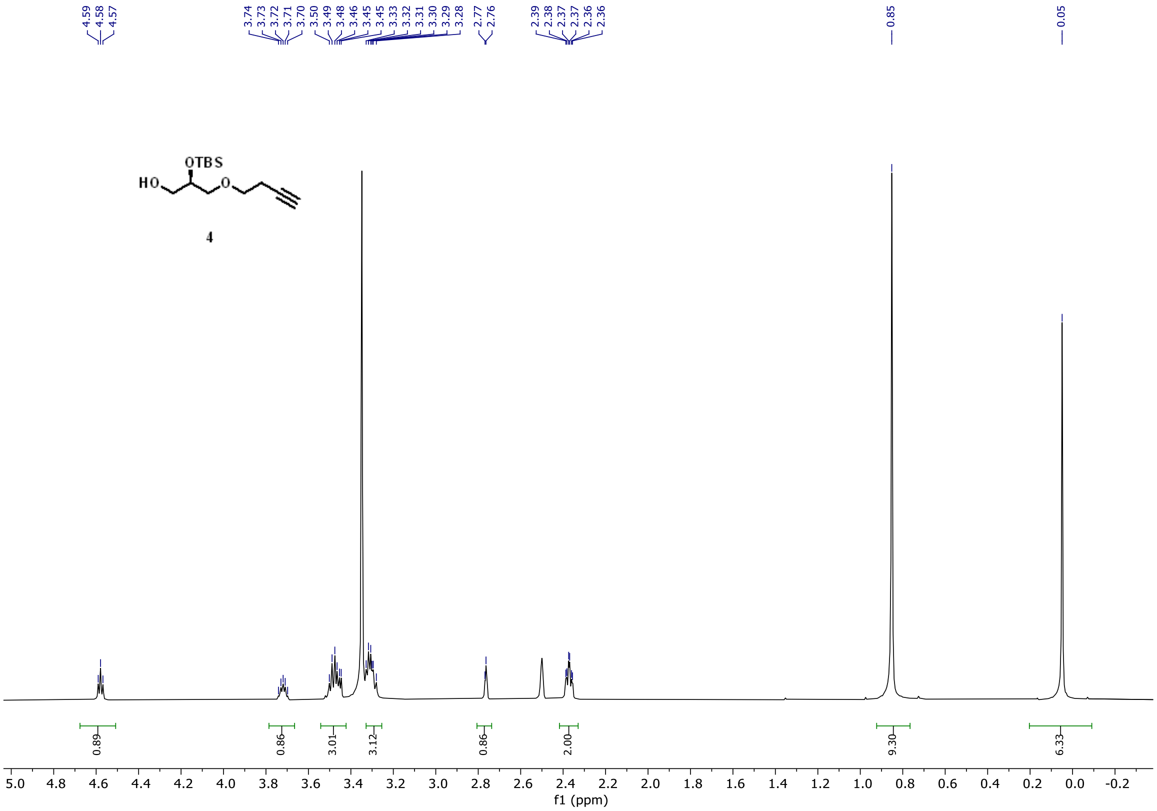

**Supplementary Figure 11**: $^{1}$H NMR Spectrum of Compound 4.

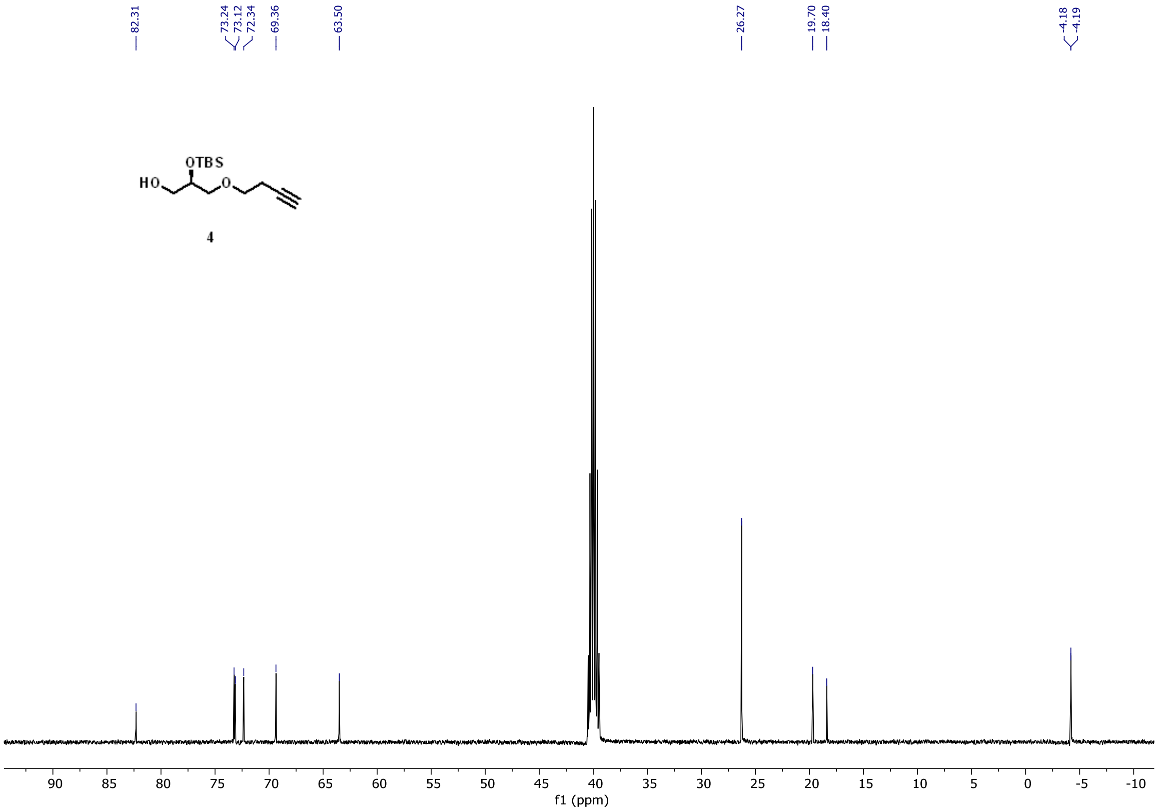

**Supplementary Figure 12**: $^{13}C$ NMR Spectrum of Compound 4.

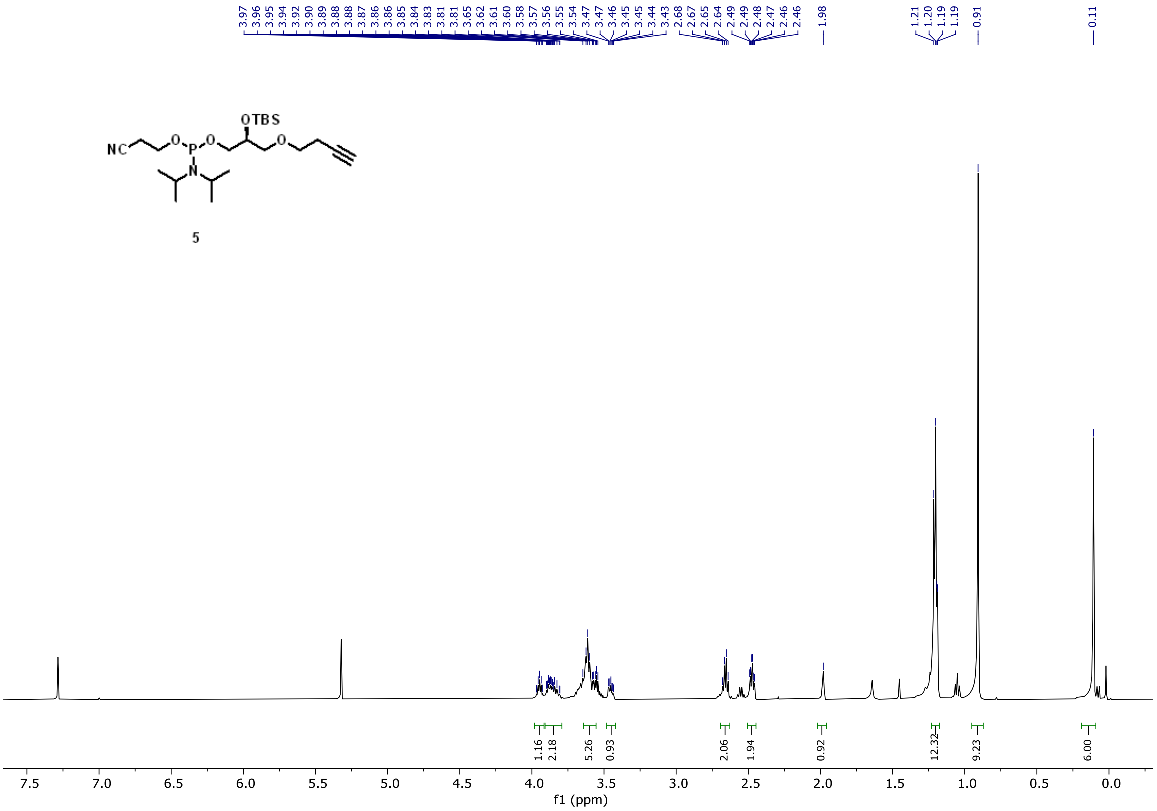

Supplementary Figure 13: $^{1}$H NMR Spectrum of Compound 5.

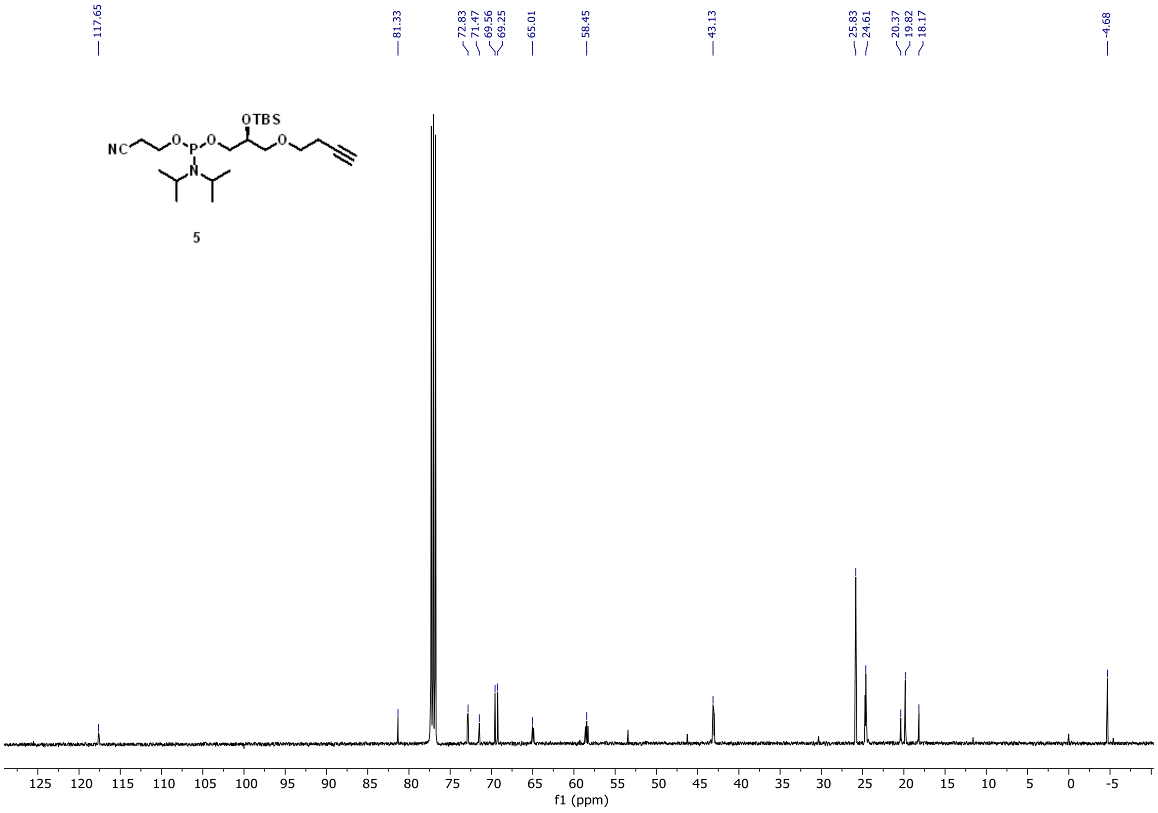

**Supplementary Figure 14**: $^{13}C$ NMR Spectrum of Compound 5.

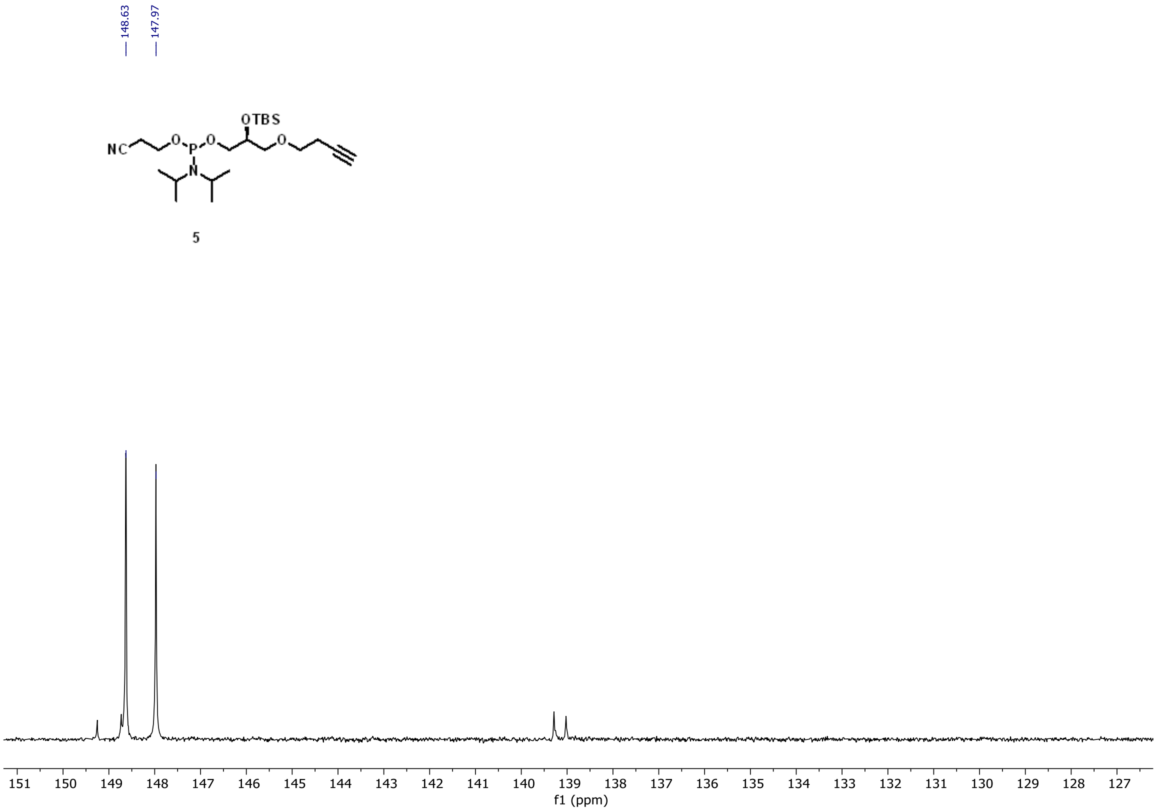

**Supplementary Figure 15**: $^{31}$P NMR Spectrum of Compound 5.

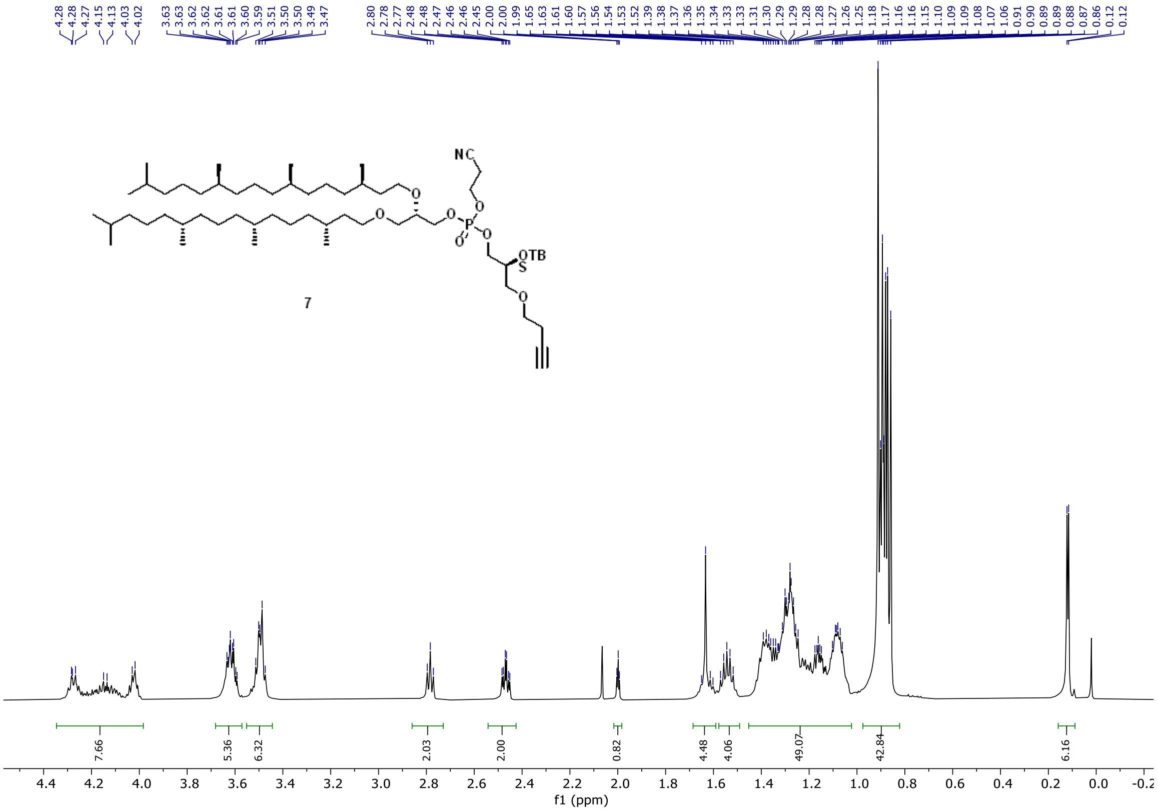

**Supplementary Figure 16**: [1]H NMR Spectrum of Compound 7.

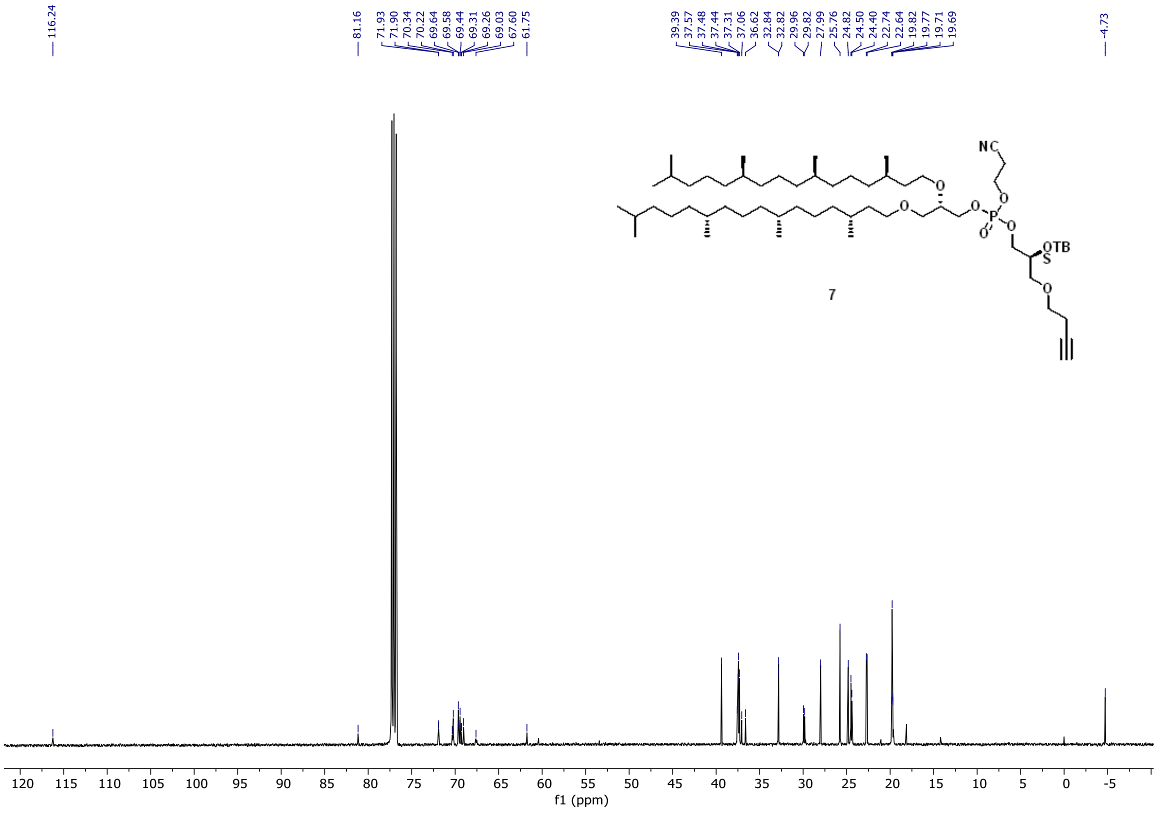

**Supplementary Figure 17**: $^{13}$C NMR Spectrum of Compound 7.

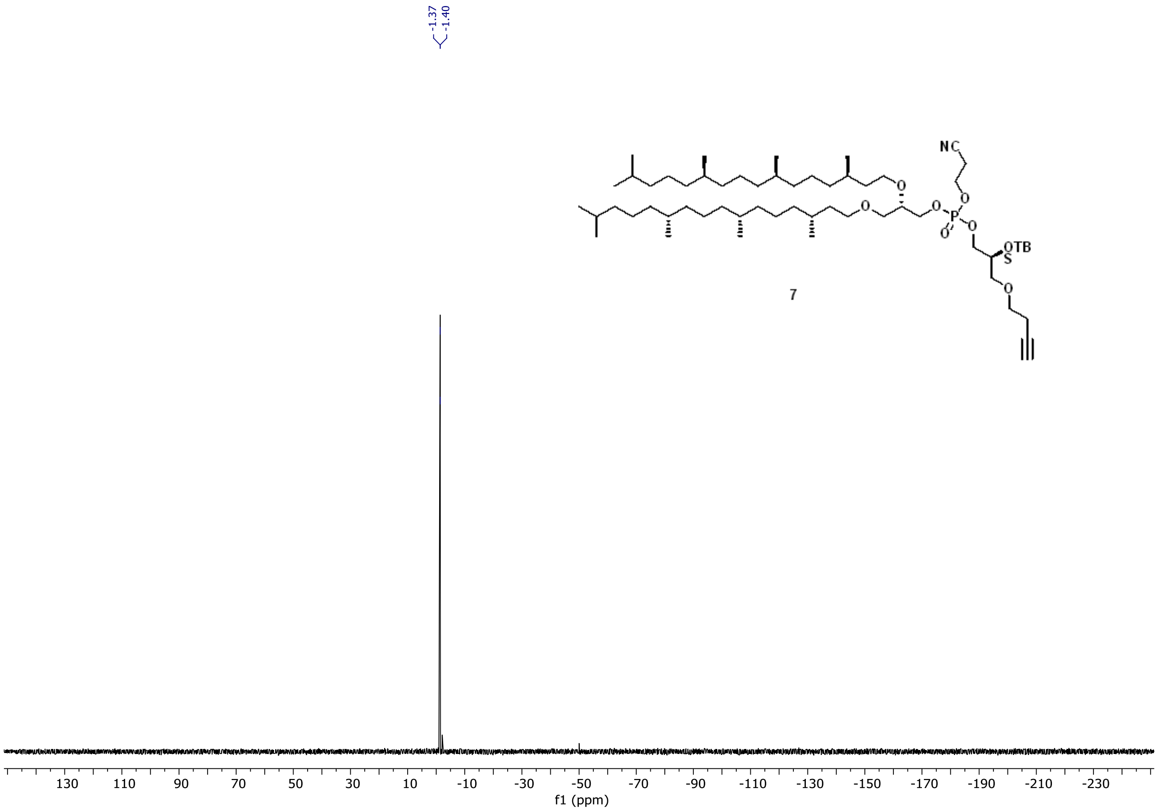

**Supplementary Figure 18**: $^{31}$P NMR Spectrum of Compound 7.

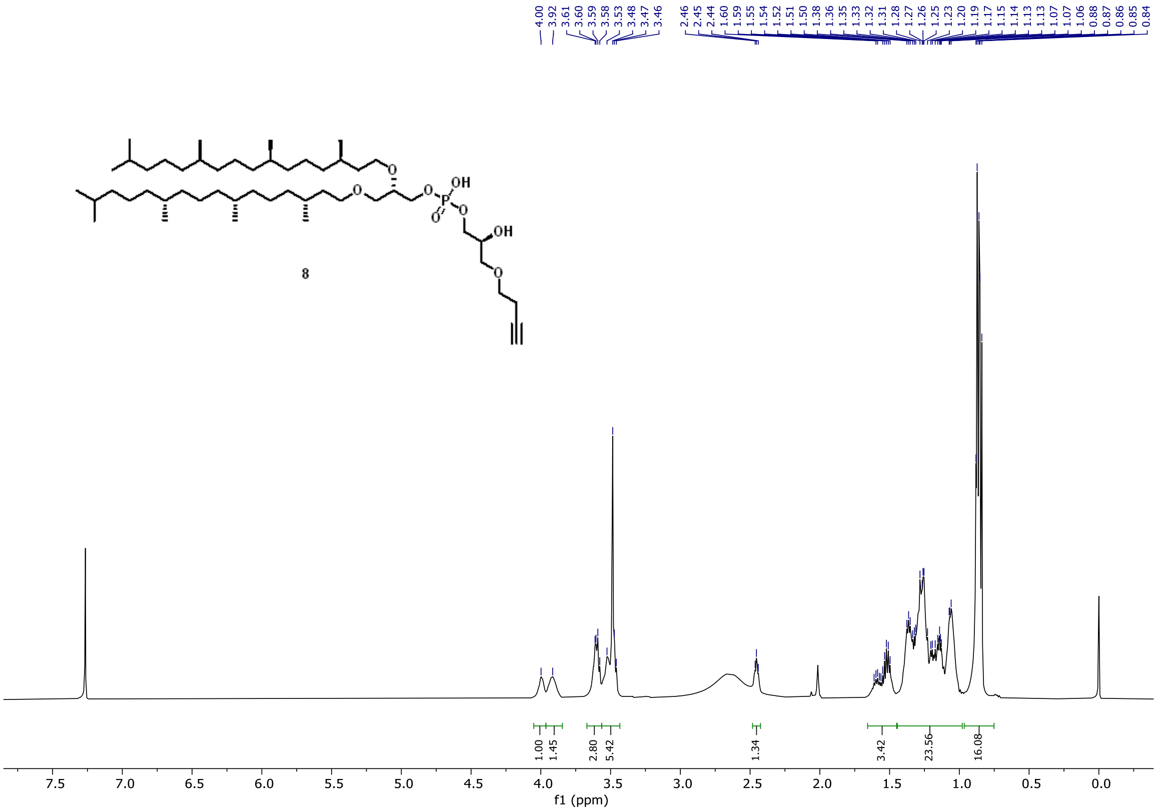

**Supplementary Figure 19**: $^{1}$H NMR Spectrum of Compound 8.

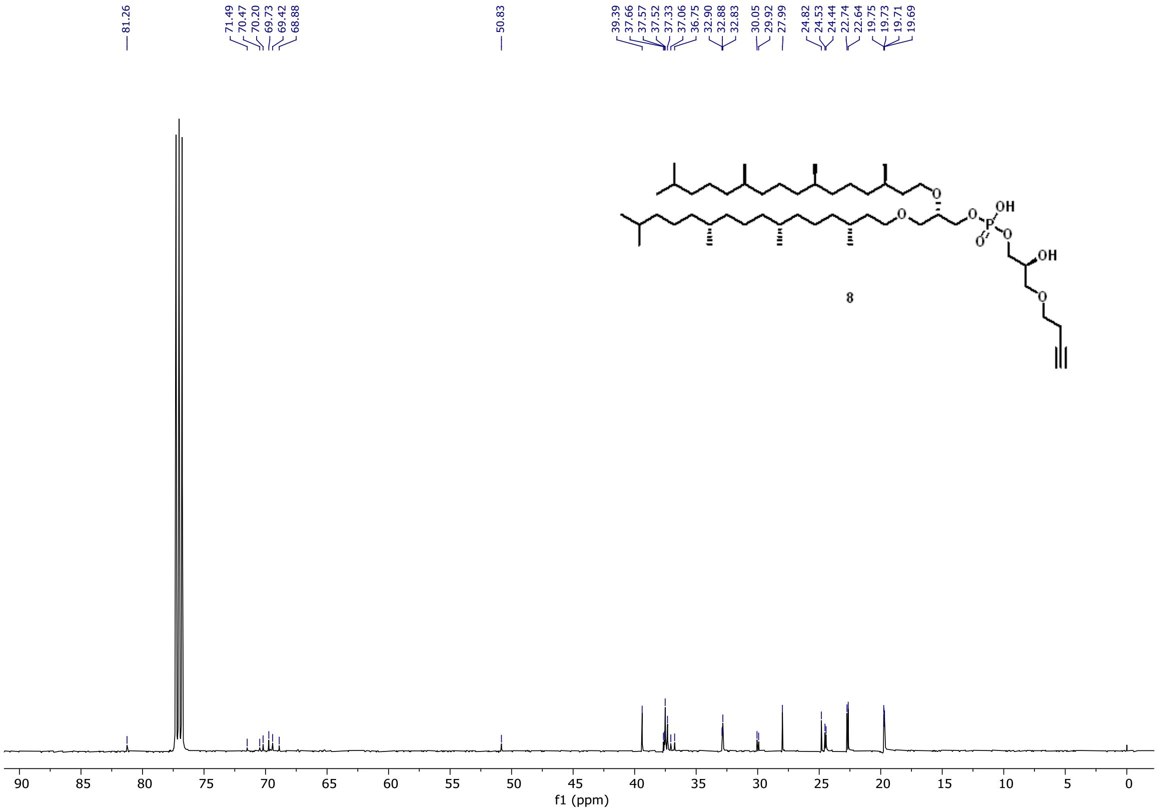

**Supplementary Figure 20**: $^{13}$C NMR Spectrum of Compound 8.

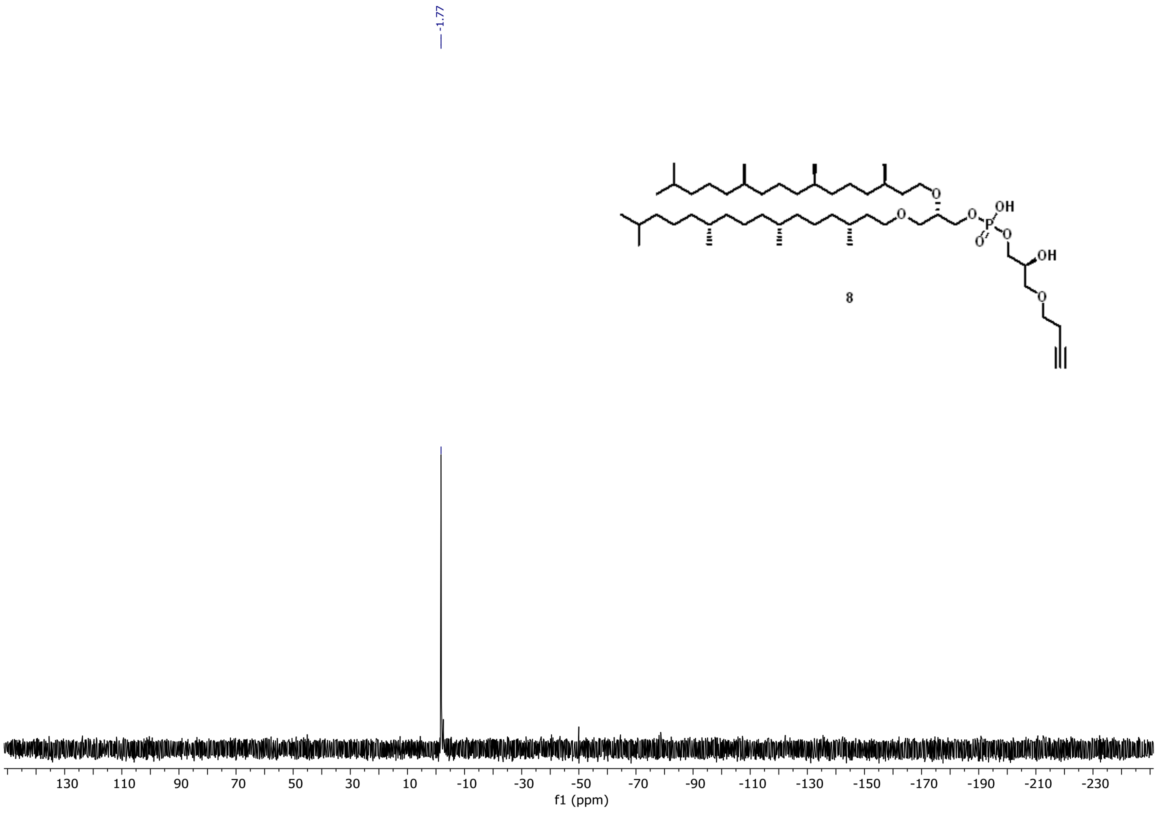

**Supplementary Figure 21**: $^{31}$P NMR Spectrum of Compound 8.

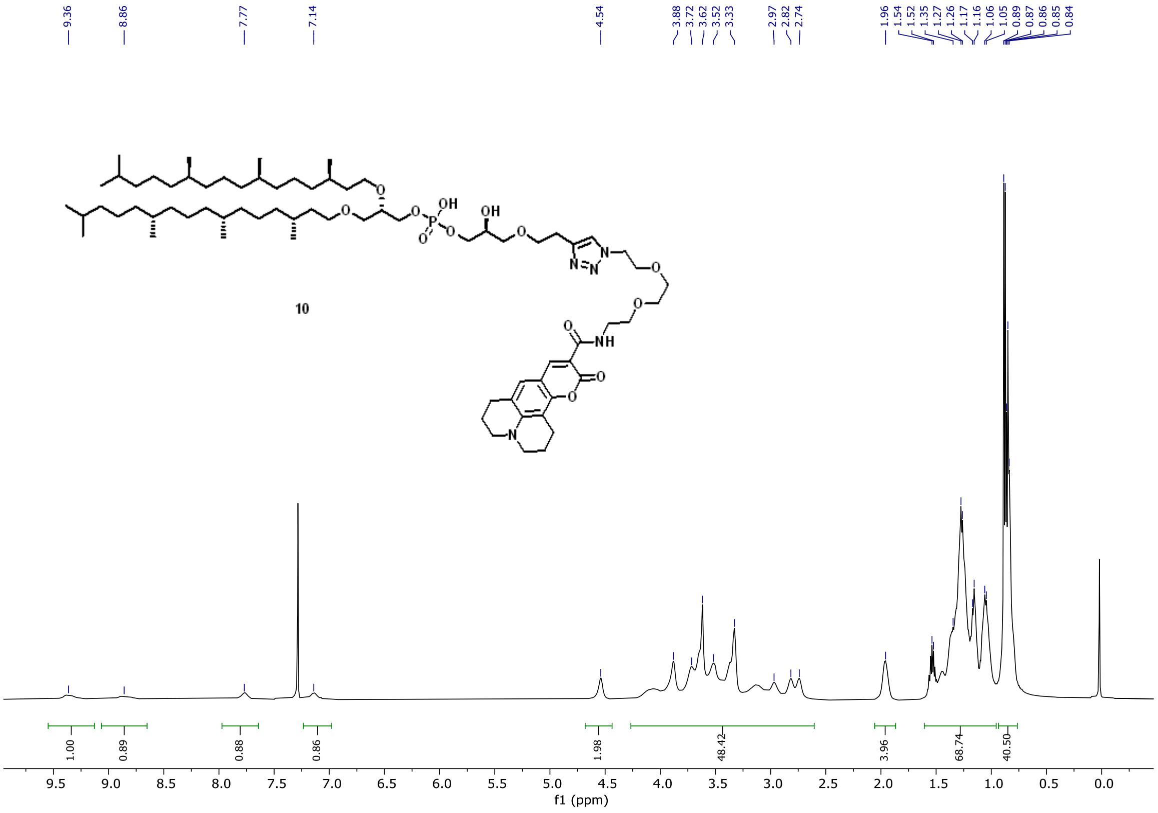

**Supplementary Figure 22**: $^{1}$H NMR Spectrum of Compound 10.

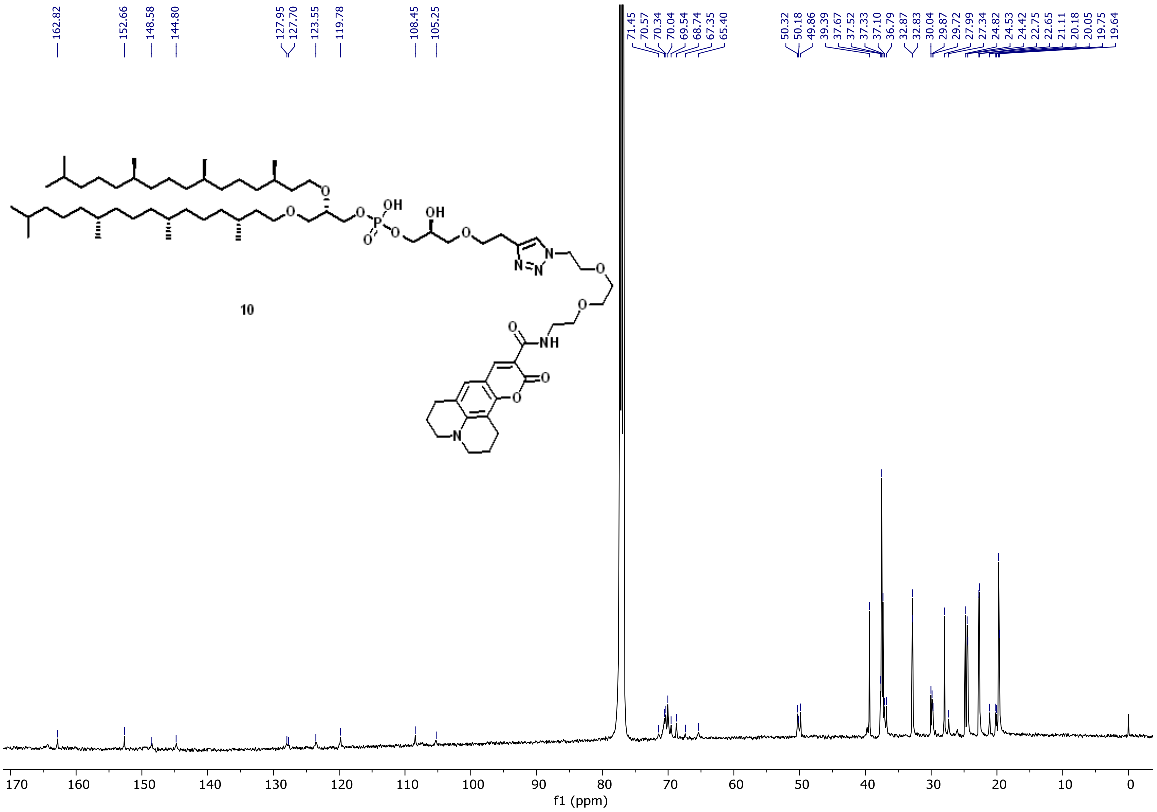

**Supplementary Figure 23**: $^{13}C$ NMR Spectrum of Compound 10.

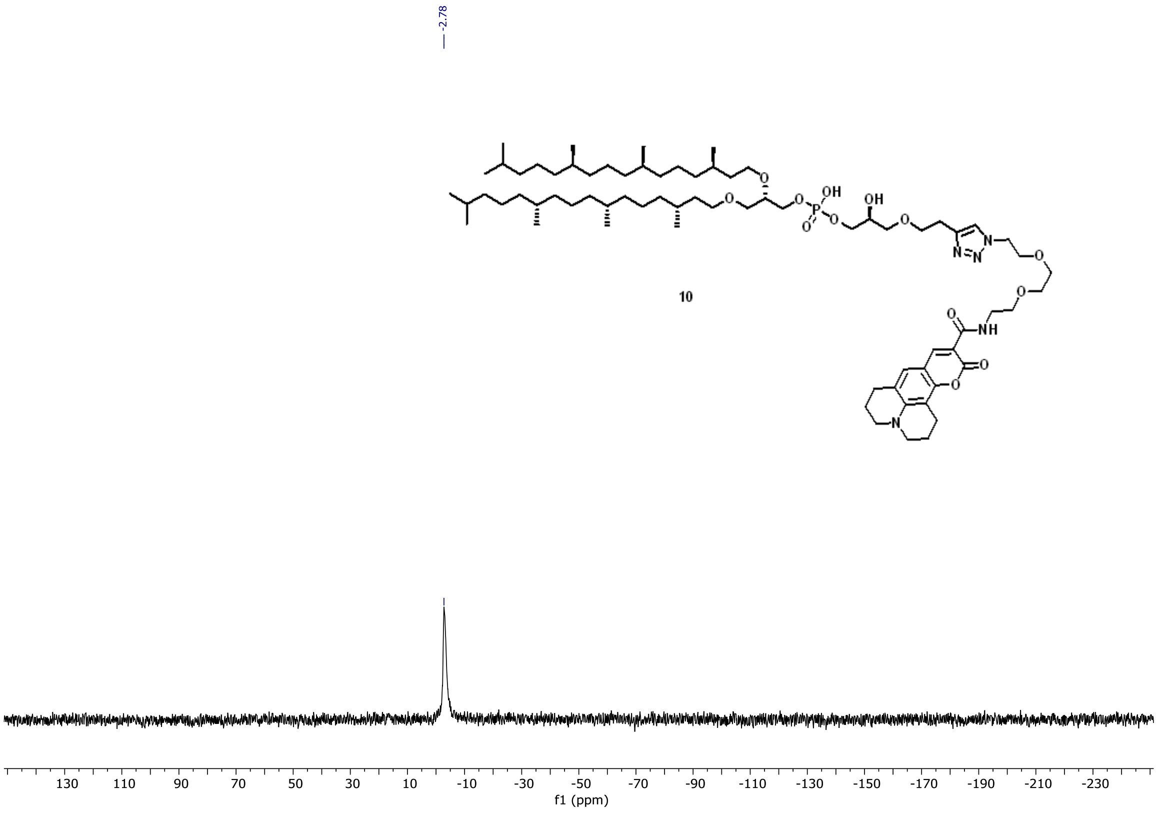

**Supplementary Figure 24**: $^{31}$P NMR Spectrum of Compound 10.

References

29 Lanz, N. D. *et al.* RlmN and AtsB as models for the overproduction and characterization of radical SAM proteins. *Methods Enzymol* **516**, 125-152 (2012). https://doi.org/10.1016/B978-0-12-394291-3.00030-7

30 Lanz, N. D. *et al.* Enhanced Solubilization of Class B Radical S-Adenosylmethionine Methylases by Improved Cobalamin Uptake in Escherichia coli. *Biochemistry* **57**, 1475-1490 (2018). https://doi.org/10.1021/acs.biochem.7b01205

31 Mocniak, L. E., Elkin, K. & Bollinger, J. M., Jr. Lifetimes of the Aglycone Substrates of Specifier Proteins, the Autonomous Iron Enzymes That Dictate the Products of the Glucosinolate-Myrosinase Defense System in Brassica Plants. *Biochemistry* **59**, 2432-2441 (2020). https://doi.org/10.1021/acs.biochem.0c00358

32 Exterkate, M. *et al.* A promiscuous archaeal cardiolipin synthase enables construction of diverse natural and unnatural phospholipids. *J Biol Chem* **296**, 100691 (2021). https://doi.org/10.1016/j.jbc.2021.100691

33 Adams, P. D. *et al.* PHENIX: a comprehensive Python-based system for macromolecular structure solution. *Acta Crystallogr D Biol Crystallogr* **66**, 213-221 (2010). https://doi.org/10.1107/S0907444909052925

34 Bunkoczi, G. *et al.* Phaser.MRage: automated molecular replacement. *Acta Crystallogr D Biol Crystallogr* **69**, 2276-2286 (2013). https://doi.org/10.1107/S0907444913022750

35 Minor, W., Cymborowski, M., Otwinowski, Z. & Chruszcz, M. HKL-3000: the integration of data reduction and structure solution--from diffraction images to an initial model in minutes. *Acta Crystallogr D Biol Crystallogr* **62**, 859-866 (2006). https://doi.org/10.1107/S0907444906019949

36 Otwinowski, Z. & Minor, W. Processing of X-ray diffraction data collected in oscillation mode. *Methods Enzymol* **276**, 307-326 (1997).

37 Emsley, P., Lohkamp, B., Scott, W. G. & Cowtan, K. Features and development of Coot. *Acta Crystallogr D Biol Crystallogr* **66**, 486-501 (2010). https://doi.org/10.1107/S0907444910007493

38 Smart, O. S., Womack, T. O., Sharff, A., Flensburg, C., Keller, P., & Paciorek, W., Vonrhein, C. and Bricogne, G. *Grade, version 1.2.20.*, <https://www.globalphasing.com.> (2011).

39 Williams, C. J. *et al.* MolProbity: More and better reference data for improved all-atom structure validation. *Protein Sci* **27**, 293-315 (2018). https://doi.org/10.1002/pro.3330

40 The PyMOL Molecular Graphics Systems v. 2.5 (Schrödinger, 2021).

41 Lloyd, C. T. *et al.* Discovery, structure and mechanism of a tetraether lipid synthase. *Nature* **609**, 197-203 (2022). https://doi.org/10.1038/s41586-022-05120-2
